# A long non-coding RNA LINC00094 regulates the transcriptional expression of lipid metabolism-related genes as a new member of core regulatory circuitry in esophageal squamous cell carcinoma

**DOI:** 10.1101/2024.07.10.602928

**Authors:** Liu Peng, Qiu-Yu Wang, Jia-Xin Chen, Yang Chen, Rong-Yao Li, Lian-Di Liao, Wan Lin, Chun-Quan Li, En-Min Li, Li-Yan Xu

**Author notes:** Corresponding authors: Dr. Li-Yan Xu, Institute of Oncologic Pathology, Shantou University Medical College, Shantou 515041, Guangdong, China. E-mail address; Dr. En-Min Li, E-mail address; Dr. Chun-Quan Li. These authors contributed equally: Liu Peng, Qiu-Yu Wang.

## Abstract

LINC00094 as a new supper-enhancer (SE)-related long non-coding RNA is associated with poor overall survival of patients with esophageal squamous cell carcinoma (ESCC). However, the transcriptional regulatory mechanism of LINC00094 and the molecular mechanisms by which LINC00094 affects the phenotype of ESCC remains unclear. Here, we found that LINC00094 promoted the proliferation of ESCC cells both in vitro and in vivo. LINC00094 knockdown significantly reduced the expression profiles of transcription activators including transcription factor 3 (TCF3) and Kruppel like factor 5 (KLF5) and lipid metabolism-related genes. Mechanically, TCF3 and KLF5 formed a core regulatory circuitry (CRC) that bound to the SEs of LINC00094 and to their own SEs to regulate the transcriptional expression in a positive feedback loop. LINC00094 recruited TCF3 and KLF5 to form a ternary complex, which forms a new CRC with TCF3 and KLF5 that regulated its own transcription as well as lipid metabolism-related genes. Knockdown of any or all three genes inhibited the expression of genes related to lipid synthesis and consistently reduced total lipid droplet levels. Treatment with SEs inhibitors (THZ1 and JQ1) effectively inhibited the formation of this CRC and the production of lipid droplets in ESCC cells. The high-risk group of CRC-associated signatures were closely associated with poor prognosis in patients with ESCC. Our findings suggest that LINC00094 is involved in the CRC by forming a complex with TCF3 and KLF5, and this regulation model can affect the phenotype of ESCC cells by controlling the expression of lipid metabolism-related genes.

**Highlights:**

1. We identified a novel functional lncRNA-LINC00094 for esophageal squamous cell carcinoma.
2. LINC00094 forms a complex with the core transcription factors TCF3 and KLF5, thereby forming a core regulatory circuitry to participate in transcriptional regulation in ESCC.
3. A core regulatory circuitry mediated by LINC00094 regulates lipid metabolism in ESCC.

## Introduction

Esophageal squamous cell carcinoma (ESCC) is one of the ten most common malignant tumors globally. More than 500,000 people die from esophageal cancer each year ^[1, 2]^. Although treatments have greatly improved in the past years, the 5-year overall survival (OS) rate of ESCC remains less than 20% ^[2]^. Therefore, it has been an especially important clinical scientific issue to determine the molecular markers for ESCC. In the past, the research on ESCC at the molecular level mainly focused on the functional proteins and their coding genes. There are still many non-coding RNAs, especially long non-coding RNAs (lncRNAs), whose functions are still undiscovered; thus, their molecular mechanisms should be explored in ESCC progression. In recent years, many upregulated lncRNAs have been identified in ESCC, which increases the degree of cancer malignancy and shortens the survival of patients ^[3–10]^. However, the mechanisms underlying such specific transcriptional regulation of the vast majority of lncRNAs and their biological significance remain uncharacterized, which extremely hinders the clinical application of lncRNAs in ESCC.

Super-enhancers (SEs) are large clusters of typical enhancers within 20kb upstream and downstream of the target molecule. SEs recruit an exceptionally large number of transcription factors (TFs) and cofactors and can be identified by extensive active histone marks, such as histone 3 lysine 27 acetylation (H3K27ac)^[11–13]^. Importantly, a large number of studies have revealed that SEs can promote many biological processes, such as tumor occurrence and development, embryonic development, immune response, and cell invasion and metastasis, by regulating key genes in these processes ^[12, 14, 15]^. Master TFs are often associated with super-enhancers themselves and form interconnected autoregulatory loops (also known as core regulatory circuitry, CRC) by binding to each other’s super-enhancers^[12, 13]^. Recent studies have suggested that some of the master TFs (KLF5, TP63, SOX2, Oct4, TCF3 or Nanog) bind to their own SEs as well as those of the other members, forming interconnected core regulatory circuitry to regulate gene expression of themselves and the other master TFs ^[12, 13, 16–20]^. Many lncRNAs that are highly expressed in cancer are regulated by SEs ^[21, 22]^. For example, co-activation of SE-driven lncRNA CCAT1 by TP63 and SOX2 promotes ESCC progression by activating the PI3K/AKT signaling pathway ^[21]^. In addition, lncRNAs can regulate the expression of master TFs and affect the activity of SEs ^[23–25]^. However, direct involvement of lncRNAs in the formation of CRC by affecting SEs activity has never been reported. Therefore, exploring the cross-regulation between lncRNAs, SEs and TFs is important for elucidating the novel functions and transcriptional regulatory mechanisms of lncRNAs as well as lncRNA-targeted cancer therapy.

In our previous study, we developed a two-stage computational approach, termed GloceRNA, to identify functional lncRNAs in ESCC ^[26]^. Moreover, we demonstrated that LINC00094 regulates the most cancer-related hallmarks and is significantly associated with ESCC prognosis. We predicted that TCF3 and KLF5 regulate LINC00094 expression by activating its SEs ^[26]^. However, whether and how TCF3 and KLF5 regulate LINC00094 transcription by affecting SE activation remains unexplored. Furthermore, it is unclear the biological function and transcriptional regulatory mechanism of LINC00094 in ESCC. Based on the above questions, in this study, we performed epigenetic profiling and transcriptomic analyses to characterize the SE landscape in ESCC cells and established ESCC dependent lncRNA-transcriptional regulatory circuitry, determined the exquisite model mediated by core lncRNA-LINC00094. Integrative analysis showed that master TFs TCF3 and KLF5 co-bind to the SE regions of LINC00094, while LINC00094 promotes TCF3 and KLF5 expression by activating their SEs. Further mechanistic exploration demonstrated that LINC00094 is used as a scaffold to recruit TCF3 and KLF5 to form a new CRC, which coordinately regulates the expression of lipid metabolism-related genes, resulting in the malignant progression of ESCC.

## Materials and Methods

### Plasmids, antibodies, shRNAs and lentiviruses

The firefly luciferase-expressing plasmids pGL3-Promoter, and Renilla luciferase-expressing plasmid pRL-TK were purchased from Promega (Madison, Wisconsin, USA). The enhancers of LINC00094, TCF3, and KLF5 were cloned into the *Kpn* Ⅰ/*Xho* Ⅰ sites of the pGL3-promoter vector. The LINC00094 enhancer region mutant plasmids were synthesized by GENEWIZ (Suzhou, China). Plasmids with mutated TCF3-and KLF5-binding motifs were constructed by replacing C with A and T with G. Lentiviral cloning vector pLKO.1-TRC and pcDNA3.1-C-Flag plasmids were purchased from Addgene (Watertown, MA, USA). The double-stranded oligonucleotide shRNAs targeting LINC00094 were cloned into the *Age* I/*Eco*R I sites of the pLKO.1-TRC lentiviral vector. Lentiviruses were generated through cotransfection of psPAX2 (Addgene) and pMD2.G (Addgene) into 293T cells using PEI transfection reagent (4 μg/mL, Sigma). After 24 h, the supernatant was collected. Puromycin at 2 μg/mL was used for selecting cells expressing shRNAs. We successfully selected the esophageal cancer cell lines KYSE150-shLINC00094 and KYSE510-shLINC00094 with stable LINC00094 knockdown. Full-length LINC00094 (NR_149319.2, 2992 nt) and LINC00094 antisense were also synthesized by GENEWIZ and cloned into the pcDNA3.1-C-Flag vector. The TCF3-and KLF5-coding sequences were cloned into the *Bam*H Ⅰ/*Xho* Ⅰ sites of the pBOBI-3×Flag vector. The clone primer sequences are provided in **Supplementary Table S1**. The siRNA and shRNA sequences are provided in **Supplementary Table S2**.

The following antibodies and reagents were used: Anti-TCF3 (Proteintech, 21242-1-AP, 1:2000 for western blotting and 8 μg for ChIP), anti-KLF5 (Santa Cruz Biotechnology, sc-398470X, 1:1000 for western blotting, and 8 μg for ChIP), anti-mouse IgG-HRP (Santa Cruz Biotechnology, sc-516102,1:5000), anti-rabbit IgG-HRP (Cell Signaling Technology, 7074s, 1:2000), normal mouse IgG (Santa Cruz Biotechnology, sc-2025, 8 μg for ChIP), normal rabbit IgG (Proteintech, B900610, 8 μg for ChIP), THZ1 (MCE, HY-80013), JQ1 (MCE, HY-13030A), TranscriptAid T7

High Yield Transcription Kit (Thermo Fisher Scientific, K0441), Pierce™ Magnetic RNA-Protein Pull-Down Kit (Thermo Fisher Scientific, 20164), HCS LipidTOX™ Green Neutral Lipid Stain (Thermo Fisher Scientific, H34475, 1:1000), Lipofectamine RNAiMAX (Invitrogen, 13778150), and Lipofectamine 3000 (Invitrogen, L3000001).

### Human clinical samples

Paired tumor and adjacent non-tumor esophageal tissues were collected from patients with ESCC following surgical resection between 2020 and 2023 at the Cancer Hospital of Shantou University Medical College. None of the patients had distant metastasis. Ethical approval was obtained from the Ethics Committees of the Medical College of Shantou University.

### Cell lines, transfection, RNA interference, and treatments

ESCC cell lines (KYSE150 and KYSE510) were established by Dr. Shimada Yutaka (Faculty of Medicine, Kyoto University, Kyoto, Japan) ^[27]^, and human embryonic kidney HEK293T cell line was obtained from the American Type Culture Collection (Manassas, VA, USA). KYSE150 and KYSE510 cell lines were cultured in RPMI 1640 medium (HYCLONE). HEK293T cell line was cultured in DMEM (Thermo Fisher). All media were supplemented with 10% fetal bovine serum (Thermo Fisher Scientific), 100 U/ml penicillin, and 100 mg/mL streptomycin. Cells were incubated at 37 °C in a humidified atmosphere containing 5% CO_2_.

The siRNA for LINC00094 was synthesized by Dharmacon (Waltham, MA, USA). The TCF3, KLF5, and negative control (NC) siRNAs were synthesized by GenePharma (Suzhou, China). siRNA transfection was performed using Lipofectamine RNAiMAX, and plasmids were transfected with Lipofectamine 3000 according to the manufacturer’s instructions. In ChIP-PCR assays, KYSE150 and KYSE510 cells were seeded into 10 cm dishes and cultured for 12–24 h until 70–80% confluence. ESCC cells were treated with THZ1 (100 nM, 12 h) and JQ1 (100 nM, 24 h), and then harvested for ChIP-qPCR assay.

### MTS assay

The MTS assay was performed as described previously ^[28]^. The transfected cells were counted and inoculated into 96-well plates with 10,000 cells per well. After 0 h, 24 h, 48 h, 72h, and 96 h of continuous culture, 20 μL MTS (Promega, G3581) was added to each well, and then incubation was continued for 2 h. The absorbance value of each pore was measured at the wavelength of 492 nm. The cell viability at different knockdown treatment groups was calculated and then mapped with GraphPad prism 7 software (San Diego, CA, USA).

### Colony formation assay

The colony formation assay was performed as described previously ^[26]^. Briefly, 1000 transfected cells per well were inoculated onto six-well plates and incubated for 14 days at 37 °C with 5% CO_2_. After washing twice with pre-cooled phosphate-buffered saline (PBS) (4 °C), cultures were fixed with ice-cold methanol and glacial acetic acid 3:1 for 15 min and stained with hematoxylin for 30 min. Colonies were photographed using ChemiDoc Touch (Bio-Rad) and the colony numbers were calculated using Image J software (US National Institutes of Health, Bethesda, MD, USA). Each experiment was performed in triplicate.

### Xenograft assays in nude mice

Five-week-old female nude mice were purchased from Vital River Laboratories (Beijing, China) for the xenograft assays. All animal studies were conducted following protocols approved by the Animal Research Committee of the Shantou Administration Center. Twelve mice were randomly separated into two groups and subcutaneously injected with 2 × 10^6^ KYSE150 cells stably transfected with shScramble and shLINC00094. Xenograft size was measured (length × width^2^) every 3 days following the appearance of tumors after injection. The measurement continued for three weeks. Mice were euthanized at the end of the experiment, and xenograft tumors were extracted for analysis.

### RNA extraction and quantitative real-time PCR (qRT-PCR)

Total RNA was extracted with TRIzol (Invitrogen) according to the manufacturer’s protocol, and 1 μg of total RNA was reverse transcribed into cDNA using HiScript III-RT SuperMix for qPCR with gDNA Eraser (Vazyme, R323-01). qRT-PCR was conducted using ChamQ Universal SYBR qPCR Master Mix (Vazyme, Q711-02) using a 7500 Real-Time PCR System (Applied Biosystems). Primers for quantitative real-time PCR are shown in **Supplementary Table S3**. *actb* expression was measured as an internal control and used for normalization.

### Western blotting and co-immunoprecipitation (co-IP)

Laemmli sample buffer (Bio-Rad) or RIPA buffer was used to lyse cells and extract total protein. Western blotting was performed for 20 μg protein using SDS-PAGE, and then transferred to PVDF membrane (Millipore). The membrane was blocked for 1 h using 5% skimmed milk powder diluted with TBST (20 mM Tris, 137 mM NaCl, 0.1% Tween-20). Membranes were incubated with primary antibodies overnight at 4 °C. The secondary antibody was incubated for 1 h at room temperature after three washes with TBST. Signals were detected using ChemiDoc Touch (Bio-Rad).

For co-IP, 1× 10^6^ cells were lysed with lysis buffer (20 mM Tris-HCl pH 7.5, 150 mM NaCl, 1 mM EDTA, 1 mM EGTA, 1% Triton X-100, 1×protease inhibitors) on ice. Exactly 500 μg of the whole-cell lysate (for each experiment) was incubated with the primary antibody or IgG on the rotary agitation overnight at 4 °C. Upon incubation with Dynabeads Protein G (Invitrogen) for 4–5 h at 4 °C, purification and western blotting were performed using indicated antibodies.

### Chromatin-immunoprecipitation (ChIP) and re-ChIP

ChIP analysis was performed as described previously ^[20, 26]^. Briefly, 2× 10^7^–2× 10^8^ cells were crosslinked with 1% formaldehyde solution (Thermo Fisher Scientific) for 10 min at room temperature and neutralized using 1.25 M glycine for 5 min, followed by two washes with cold PBS. These cells were lysed twice with 1 mL cell lysis/wash buffer (150 mM NaCl, 0.5 M EDTA pH 7.5, 1 M Tris pH 7.5, 0.5% NP-40, 1% Triton X-100, 1×protease inhibitors). The cells were mechanically lysed by pipetting up and down several times with an insulin syringe. Cell pellets were resuspended in 1 mL of shearing buffer (1% SDS, 10 mM EDTA pH 8.0, 50 nM Tris pH 8.0, 1×protease inhibitors) and sonicated (Covaris E220) to release 100–500 bp fragments. The supernatants were then diluted using the dilution buffer (0.01% SDS, 1% Triton X-100, 1.2 mM EDTA pH 8.0, 150 nM NaCl). Subsequently, anti-KLF5, anti-TCF3, or normal IgG was added to each sonicated chromatin sample and incubated at 4 °C overnight. Then, these complexes were conjugated to Dynabeads Protein A/G magnetic beads (Invitrogen) for 4–6 h at 4 °C. Dynabeads were washed five times with cold wash buffer and once with cold TE buffer. Next, the DNA fragments were eluted using freshly prepared elution buffer (130 mM NaHCO_3_, 1% SDS). DNA molecules were vortexed for 15 min and exposed to reverse crosslinking using 5 M NaCl at 65 °C overnight. The next day, DNA samples were treated with RNase A and proteinase K and purified using the QIAquick PCR purification kit (QIAGEN). The purified DNA was analyzed via qRT-PCR or deep sequencing.

For tissue ChIP-seq, 50-100 mg fresh cancer and normal tissues were dissected and snap-frozen in liquid nitrogen for each ChIP assay. The tissue fragments frozen in liquid nitrogen were added to 1% formaldehyde/DMEM buffer and ground four times on ice using a Dounce Tissue Grinder. The tissue suspension was filtered through a 5 mL Round Bottom Polystyrene Test Tube with Cell Strainer (BD Flacon, 352235), and the cell suspension obtained by filtration was fixed in 1% formaldehyde/DMEM buffer for 30 min at room temperature. Glycine was added at a final concentration of 125 mM to terminate the fixation. The following steps were consistent with the cell ChIP experiment.

For re-ChIP assay^[29]^, protein-DNA-bead complexes were washed thrice with re-ChIP wash buffer (2 mM EDTA, 500 mM NaCl, 0.1% SDS, 1% NP40), followed by a double wash with 1× TE buffer. The washed immunoprecipitated protein-DNA complexes were eluted via incubation for 30 min at 37°C in 75 μL of re-ChIP elution buffer (1× TE, 2% SDS, 15 mM DTT, 1×protease inhibitor) and diluted 1:50 in ChIP dilution buffer (with 50 μg of bovine serum albumin and protease inhibitor) followed by re-IP with the secondary antibodies. Samples were analyzed using ChIP-PCR with primers listed in **Supplementary Table S4**.

### ChIP-seq data analysis

H3K27ac ChIP-seq data for seven ESCC cell lines KYSE70, KYSE140, KYSE150, KYSE510, TT, TE5, and TE7 were generated from our previous studies using the Gene Expression Omnibus (GEO) datasets (GSE106563, GSE131493, and GSE106434) ^[13, 21, 22]^. ChIP-seq data of TCF3 and KLF5 were generated in both KYSE150 and KYSE510 cell lines and two ESCC and normal tissues pairs. ChIP-seq data were processed as described previously ^[26]^. In brief, sequencing reads were aligned to the reference genome (GRCh38) with Bowtie Aligner (v0.12.9). Peaks were identified with MACS2 (v 2.1.1). Wiggle files were generated and normalized at the unit of reads per million reads. We converted wiggle files into bigwig files using the WIGTOBIGWIG tool (http://hgdownload.cse.ucsc.edu/admin/exe/) and then visualized them using Integrative Genomics Viewer (IGV, http://www.broadinstitute.org/igv/home). The ROSE method defined enhancers as H3K27ac peaks that are 2 kb away from any TSS. Following enhancer element stitching within 20 kb both upstream and downstream of the target gene, typical enhancers and SEs were then classified using a cutoff at the inflection point (tangent slope=1) based on the ranking order. ROSE GeneMapper.py method was used to identify the genes regulated by the SE. Both SEs and TEs were assigned to the overlap, proximal, and closest genes to the center of the stitched enhancer. If genes appeared in the overlap, proximal, or closest SEs, they were considered SE-associated genes.

### Transcriptome sequencing (RNA-seq) and data analysis

The RNA-seq data of siLINC00094, siTCF3, and siKLF5 were generated in KYSE150 and KYSEE510 cell lines. First, RNA-seq reads were aligned to the human reference genome (GRCh38) using the hisat2 software. Second, gene expression was quantified using the Stringtie software. Finally, DESeq2 was used to identify the differentially expressed genes (DEGs) (*P*<0.05, Fold change>1.2) based on the read counts. Gene annotation and analysis were conducted using metascape (http://metascape.org/) and OmicShare (https://www.omicshare.com/).

### Luciferase reporter assay

KYSE150 and KYSE510 cells were seeded onto 96-well plates. Cells were co-transfected with a reporter firefly luciferase-expressing pGL3-promoter-enhancer plasmid (1 μg), and a Renilla luciferase-expressing plasmid pRL-TK (20 ng) as a normalization control. In the TCF3/KLF5/LINC00094 knockdown and TCF3/KLF5/LINC00094 overexpression experiments, ESCC cells were first transfected with TCF3/KLF5/LINC00094 siRNAs or TCF3/KLF5/LINC00094 plasmid, respectively. After 48 h of transfection, the luciferase activity was measured using the Dual-Luciferase Reporter Assay System (Promega).

### RNA-binding protein immunoprecipitation (RIP) assay

RIP assay was performed as described previously ^[6]^. Before the RIP assay, cells on ice in an open dish were placed in a Stratalinker UV-light box and irradiated for 2 min. After UV crosslinking, cells were washed with pre-cooled PBS and softly scraped. Cell pellets were lysed in RIP buffer (50 mM Tris-HCl pH 7.5, 150 mM NaCl, 5 mM EDTA, 0.5% NP40, 1% Triton X-100, 1% RNase inhibitor) for 15 min at 4 °C, shaking. Cell pellets were lysed again with a small volume of RIP buffer. Then RNA from 10% of the lysate was extracted using TRIzol (Invitrogen) to serve as the ‘input’. The remainder of the cell lysate was first incubated with Dynabeads Protein A/G to remove non-specific binding. The precleared lysate was incubated with anti-TCF3, anti-KLF5, and normal IgG antibodies at 4 °C overnight. Then, the RNA-protein complexes were conjugated to Dynabeads Protein A/G magnetic beads for 4–6 h at 4 °C. After beads were sequentially washed 4–6 times in RIP buffer, RNA was isolated using TRIzol, incubated with DNase I (NEB) and subjected to qRT-PCR.

### RNA-pull down assay

Biotin-labeled RNAs were synthesized using Scientific TranscriptAid T7 High Yield Transcription Kit (Thermo Fisher Scientific). Biotin-labeled RNA-pulldown assay was performed using Pierce™ Magnetic RNA-Protein Pull-Down Kit (Thermo Fisher Scientific). Briefly, LINC00094 was cloned into the pcDNA3.1 vector with the T7 promoter and then used to amplify DNA templates for RNA synthesis. RNA was transcribed in vitro using T7 RNA polymerase and biotin labeling mix, treated with RNase-free DNase I, and purified with the GeneJET RNA Purification Kit (Thermo Fisher Scientific). Cell lysates were incubated with biotin-labeled RNAs overnight. Proteins associated with biotin-labeled RNAs were immunoprecipitated with streptavidin magnetic beads and subjected to western blotting analysis.

### Cell fractionation assay

Cell fractionation assay was performed as described previously ^[30]^. KYSE150 and KYSE510 cell fractions enriched for chromatin were obtained. Exactly 2× 10^7^ cells were washed with pre-cooled PBS and lysed with 1 mL Buffer A (10 mM Tris-HCl pH 7.5, 3 mM CaCl_2_, 3 mM MgCl_2_, 1 nM DTT, 320 mM Sucrose, 0.3% NP40, 1×protease inhibitors, 1% RNase inhibitor) on ice. Cells were centrifuged for 5 min at 2800 ×g after being lysed for 10 min. The supernatant was collected in another centrifuge tube; 100 μL aliquots were removed and mixed with SDS loading buffer. RNA was extracted from the remainder of the cell supernatant using TRIzol. This component served as the cytoplasm RNA. Crude nuclei were washed in Buffer A twice, before adding two packed cell volumes of Buffer B (1.5 mM MgCl_2_, 420 mM NaCl, 200 mM EDTA, 25% glycerol, 1 mM DTT, 20 mM HEPES pH 7.7, 1×protease inhibitors, 1% RNase inhibitor). Nuclei were extracted for 30 min at 4 °C and centrifuged for 15 min at 8000 ×g. Half of this supernatant was mixed with SDS loading buffer, and RNA was extracted from the remainder using TRIzol. This component served as the nucleoplasm RNA. Half of the cell pellets were lysed in 1×SDS buffer, and RNA was extracted from the remainder of pellets using TRIzol. This component served as the chromatin RNA.

### Immunofluorescence

KYSE150 and KYSE510 cells were seeded onto coverslips and fixed with 4% paraformaldehyde for 10 min at room temperature. After washing with PBS, the cells were permeabilized with 0.1% Triton X-100 for 8 min on ice. Then, cells were stained with HCS LipidTOX™ Green Neutral Lipid Stain (1:1000) for 1 h at room temperature. After washing, cells were analyzed using the Zeiss LSM800 confocal microscope.

### Least shrinkage and selection operator (LASSO) Cox regression algorithm

Non-coding RNA expression data (GSE53625, PRJNA665149) of ESCC were retrieved from the GEO database (https://www.ncbi.nlm.nih.gov/geo/query/acc.cgi?acc=GSE53625) and SRA database (https://www.ncbi.nlm.nih.gov/bioproject/PRJNA665149)^[31]^. The LASSO Cox regression algorithm is a variation of LASSO and was applied to prioritize the most relevant prognostic candidates of a DEGs set, based on which the regression risk model was established and applied to the clinical data of the training cohort (GSE53625, n=179) and independent validation cohort (TCGA, n=84). First, 3781 DEGs (FC >1.2) were selected from the RNA-seq data of LINC00094 knockdown in KYSE150 cells. Second, univariate Cox regression analyses were conducted to obtain 212 independent prognostic genes significantly correlated with OS in 179 patients with ESCC. Third, LASSO Cox regression analysis was performed to further select candidate genes significantly related to ESCC and the 10-fold cross-validation was used to determine the optimal lambda value, limiting the errors to a minimum of one standard error. Finally, we built a 28-gene signature based on the association between DEGs upon LINC00094 knockdown and OS in the training cohort (n = 179). Two cohorts of patients with ESCC were separated into high-and low-risk groups using the median value as the cutoff. Kaplan-Meier analysis and receiver operating characteristic curve (ROC) analyses were performed to evaluate the accuracy and sensitivity of the model using the R package “survival” and “survivalROC”, respectively.

### Statistical analysis

The analyses were performed using SPSS software (ver.13.0) or R 3.1.2 for windows. Kaplan-Meier curve was constructed for OS analysis using a Log-rank test. Differences between groups were evaluated using the Student t-test. *P* < 0.05 was considered to be statistically significant. Graphs were established using GraphPad Prism software, and data were shown as the mean ± SD (standard deviation) or SEM (standard error of the mean).

## Results

### LINC00094 promotes malignant phenotypes of ESCC cells

Little is known regarding the regulation or function of LINC00094 in human cells. The public dataset (GSE53625, n=179) validated the up-regulation of LINC00094 in ESCC samples compare with matched nonmalignant esophageal mucosa (**Figure 1A**). Meanwhile, we detected the expression of LINC00094 in normal esophageal epithelial cells and ESCC cells by qRT-PCR. The results showed that the expression of LINC00094 in ESCC cells, especially KYSE150 and KYSE510 cells, was significantly higher than that in normal esophageal epithelial cells (NE1, NE2, NECA6) (Supplementary Figure 1B). Notably, high-expression of LINC00094 was significantly associated with poor overall survival of ESCC patients (*P*=0.0341, **Figure 1B**), indicating the biological significance of LINC00094. Moreover, we first confirmed that knockdown of LINC00094 greatly reduced proliferation and clonogenicity of ESCC cells (**Figure 1C-F**). Subsequently, the KYSE150 cells stably transfected with shScramble or shLINC00094 were inoculated subcutaneously in nude mice. The xenograft data showed that LINC00094 knockdown resulted in a marked reduction in mass of the tumors (**Figure 1G-I**). These data identified LINC00094 as a functionally oncogenic lncRNA in ESCC.

**Figure 1.**
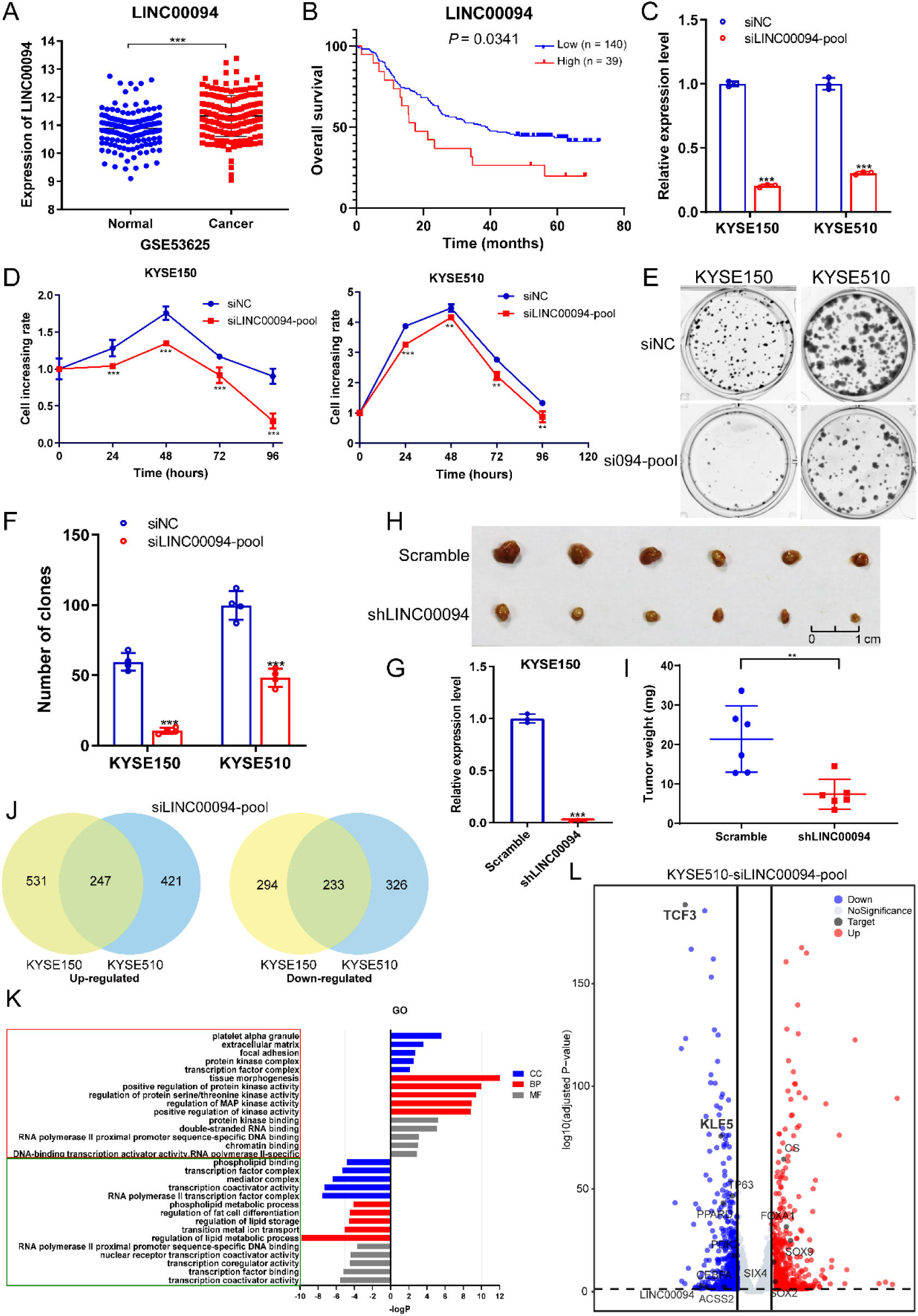
LINC00094 regulates cell proliferation and lipid metabolism pathway in ESCC. **A.** mRNA levels of LINC000094 were derived from the indicated GEO datasets measuring matched ESCC and adjacent non-malignant esophageal epithelium. **B.** Kaplan-Meier plot showing the association between LINC00094 expression and overall survival in ESCC patients. **C.** qRT-PCR analysis validating the silencing of LINC00094 in KYSE150 and KYSE510 cells. **D.** MTS assay after LINC00094 knockdown in ESCC cells. **E-F.** LINC00094 knockdown using individual siRNA-pool inhibited cell proliferation in colony formation assay. E representative images (si094-pool: siLINC00094-pool); F, quantitative analyses of colony numbers. The mean ± SD is shown n≥3. * *P* < 0.05, ** *P* < 0.01, *** *P* < 0.001. *P*-values were determined using a two-sided t-test. **G.** LINC00094 was stably silenced using shRNA in KYSE150 cells. **H-I.** Mouse xenograft assay upon LINC00094 knockdown using shRNA via intraperitoneal injection. H, images of dissected tumors; I, the mean tumor weight measured at the end point. The mean ± SEM are shown, n = 6 (the numbers of tumor in one group), ** *P* < 0.01. *P*-values were determined using a one-sided t-test. **J.** KYSE150 and KYSE510 cell differential gene venny map after knocking down LINC00094 (FC>1.5, *P*<0.05). Left, high DEGs; Right: low DEGs. **K.** The biological processes associated with genes affected by LINC00094 knockdown. The red box represents the enrichment pathways of high DEGs, and the green box represents the enrichment pathways of low DEGs. **L.** After LINC00094 knockdown in KYSE510 cells, the volcano plot of DEGs, red represents the differentially upregulated genes and blue represents the differentially downregulated genes.

Subcellular fractionation was first performed to characterize the mechanisms underlying LINC00094-mediated cellular effects, which identified the localization of LINC00094 in ESCC cells. qRT-PCR of fractionated nuclear and cytoplasmic RNA confirmed LINC00094 predominant presence in the nucleus (Supplementary Figure 1B). Subsequently, to further explore the function of LINC00094 in ESCC cells, RNA-seq was performed upon silencing of LINC00094 in KYSE150 and KYSE510 cells. Merged with RNA-seq data, we identified a total of 233 differentially downregulated transcripts and 247 differentially upregulated transcripts (*P*<0.05, FC>1.5) (**Figure 1J**). Gene Ontology (GO) analysis showed that downregulated genes upon LINC00094 silencing in ESCC cells were strongly enriched for transcriptional regulation-related phenotypes and lipid metabolism-related phenotypes, including transcription regulator and coregulator activity, transcription factor complex, lipid and phospholipid metabolic process and lipid storage (**Figure 1K**). Volcano plot analysis showed that the expression of many core transcription factors and lipid metabolism-related enzymes were altered after knockdown of LINC00094 (**Figure 1L, Supplementary Figure 2**). Altogether, LINC000094 promotes the malignant phenotype of ESCC by regulating the gene transcription.

### LINC00094 regulates the transcriptional expression of lipid metabolism-related genes

We hypothesized that LINC00094 is located in the nucleus and may be involved in the transcriptional regulation, thus exerting its biological functions such as regulating lipid metabolic processes. Volcano plot analysis was performed to explore this hypothesis, revealing that LINC00094 silencing led to abnormal expression of many TFs, such as TP63, TCF3, KLF5, PPARD and CEBPA, most of which are associated with the lipid metabolism process (**Figure 1K, Supplementary Figure 2**)^[20, 32–35]^. Knocking down LINC00094 led to a significant reduction in TCF3 and KLF5 expression, which caught our attention. Indeed, we further verified that silencing LINC00094 with siRNAs in ESCC cells markedly decreased expression of TCF3 and KLF5 at both mRNA and protein levels (**Figure 2A**), while LINC00094 overexpression produced the opposite effect (**Figure 2B**). Overexpression of LINC00094 in ESCC cells after knocking down TCF3 and KLF5 also showed that LINC00094 could effectively rescue the reduced expression caused by silencing TCF3 and KLF5 (**Figure 2C**). RNA extracted from tumors of xenografted mice and qRT-PCR assay showed that the expression of TCF3 and KLF5 was also significantly reduced in the LINC00094 knockdown group (**Figure 2D**). Furthermore, to verify whether LINC00094, TCF3 and KLF5 are involved in regulating lipid metabolism in ESCC, we detected lipid droplet formation after knockdown of the three molecules and found that lipid droplet synthesis was significantly reduced in KYSE150 and KYSE510 cells (**Figure 2E-J**). These data showed that LINC00094 regulates the expression of TCF3 and KLF5 and that all three molecules modulate lipid droplet formation in ESCC cells.

**Figure 2.**
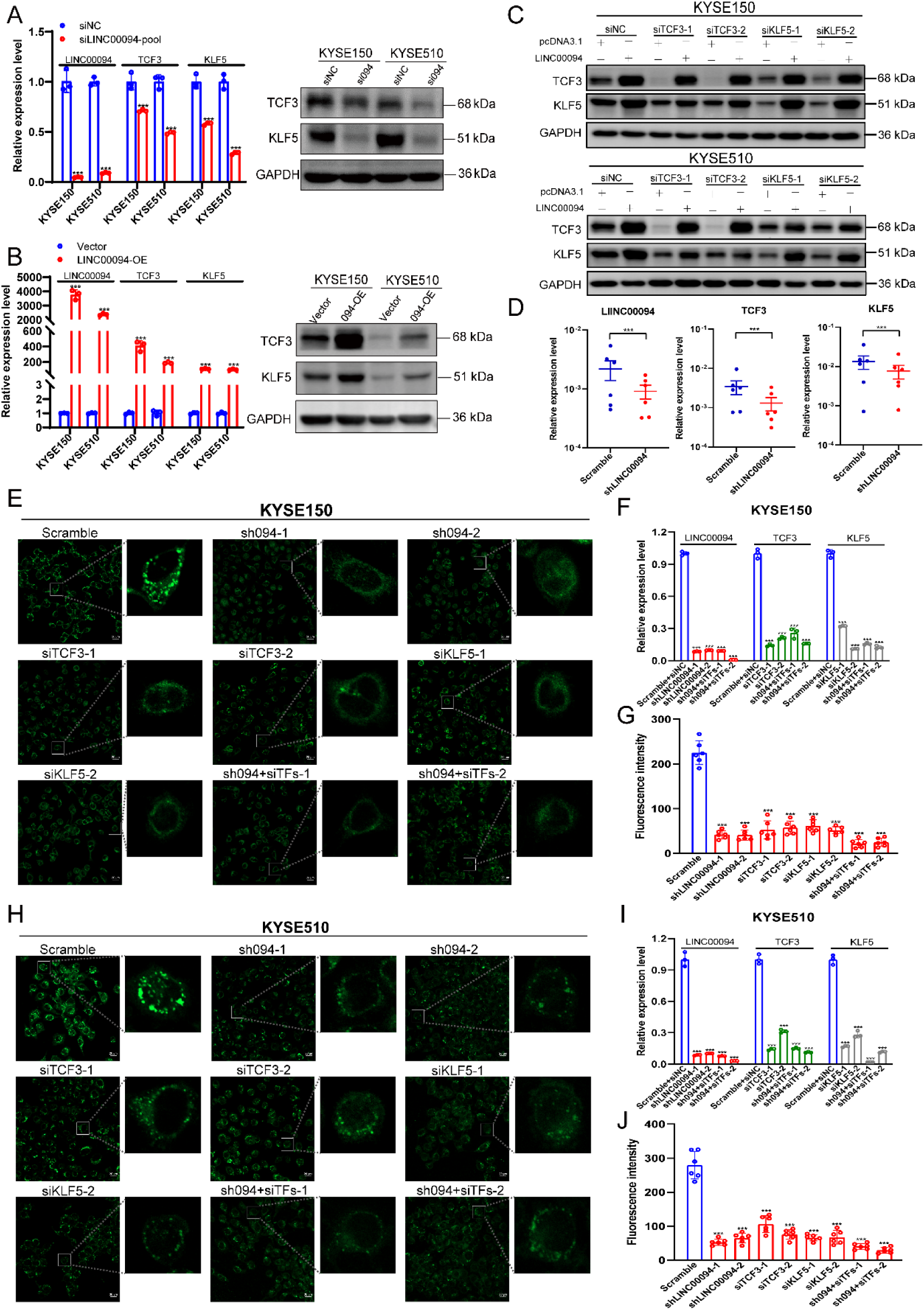
LINC00094 promotes TCF3 and KLF5 expression as well as lipid droplet formation. **A.** Left, relative mRNA levels of TCF3 and KLF5 following siRNA knockdown of LINC00094 in KYSE150 and KYSE510; right, western blotting detection of TCF3 and KLF5 in two ESCC cell lines upon silencing of LINC00094 (si094: siLINC00094). **B.** Left, relative mRNA levels of TCF3 and KLF5 after overexpressing LINC00094; right, western blotting detection of TCF3 and KLF5 in two ESCC cell lines upon overexpressing LINC00094 (094-OE: LINC00094 overexpression). **C.** Western blotting analyses showing protein levels of TCF3 and KLF5 upon either knockdown of TCF3 and KLF5 alone or combined with full-length LINC00094 overexpression in KYSE150 and KYSE510 cells. **D.** Relative mRNA levels of LINC00094, TCF3, and KLF5 after shRNA knockdown of LINC00094 in xenograft mouse tumors. The mean ± SEM are shown, n = 6, *** *P* < 0.001. *P*-values were determined using a two-sided t-test. **E.** Confocal images of lipid droplets after LINC00094, TCF3, or KLF5 knockdown alone or together in KYSE150 cells as well as qRT-PCR assay for different treatment groups (**F**) (sh094: shLINC00094; siTFs: siTCF3+siKLF5). Scale bar, 20 μm. **G.** Quantitative analysis of lipid droplet staining based on the confocal images. Mean ± SD are shown, n = 6, as the number of microscopic vision. * *P* < 0.05, ** *P* < 0.01, *** *P* < 0.001. *P*-values were determined using a two-sided t-test. **H-J.** Confocal images of lipid droplets and quantitative analysis in KYSE510 cell.

### The expression of LINC00094 was regulated by TCF3 and KLF5 through a positive feedback loop

In our previous study, we predicted that the core transcription factors TCF3 and KLF5 regulated the expression of LINC00094 through SEs ^[26]^. However, the mechanism of TCF3 and KLF5 regulating LINC00094 transcription remains unexplored. To explore how TCF3 and KLF5 regulated LINC00094, two super-enhancer inhibitors, THZ1 and JQ1, were used first. THZ1 is a CDK7 inhibitor, the THZ1-sensitive transcripts demonstrated that they were frequently associated with SE, consistent with its known function ^[36, 37]^. JQ1 is a well-known bromodomain inhibitor with a high level of potency against BRD4, and it has been shown to effectively reduce the expression of SE-related genes ^[37–39]^. Importantly, we observed decreased expression of LINC00094, TCF3, and KLF5 in a dose-dependent manner upon THZ1 or JQ1 treatment in ESCC cells (Supplementary Figure 3). Moreover, we analyzed H3K27ac ChIP-seq profiles in seven ESCC cells. The SE region of LINC00094 was found in all seven ESCC cell lines, including three typical enhancer elements (E1, E2 and E3) (Supplementary Figure 4A) ^[20, 26]^. Notably, TCF3 and KLF5 ChIP-seq data from KYSE150 and KYSE510 cells and paired tumor and normal tissues of patients with ESCC revealed multiple binding peaks consistent within the LINC00094 SEs (Supplementary Figure 4A). The other observation was that the SE flanking LINC00094 was ESCC tissue-specific since they were either undetectable or much weaker in normal tissues (Supplementary Figure 4A). Consistently, motif analysis revealed that the binding motifs of TCF3 and KLF5 were enriched in the SE of LINC00094 (Supplementary Figure 4A). ChIP-qPCR was performed to validate the ChIP-seq results. As anticipated, ChIP-qPCR confirmed the enrichment of TCF3 and KLF5 at SEs (E1, E2, and E3) of LINC00094, THZ1 and JQ1 inhibited the interaction of TCF3 and KLF5 with the LINC00094 SE (Supplementary Figure 4B-C). Indeed, we validated that either TCF3 or KLF5 knockdown caused significant reduction of the mRNA level of LINC00094, while overexpression either of TFs induced LINC00094 expression in ESCC cell lines (Supplementary Figure 4D-E). Subsequently, to measure the direct regulatory effect of TCF3/KLF5 on its target enhancer elements, we further performed site-directed mutagenesis to mutate TCF3 and KLF5 binding motifs on E1, E2, and E3 elements of LINC00094. Additionally, overexpression of TCF3 or KLF5 increased the luciferase reporter activity of the wild-type enhancer significantly, and TCF3 or KLF5 knockdown produced the opposite effects but produced no detectable effect on the mutant enhancer (Supplementary Figure 5). These data confirm the direct regulation of both TCF3 and KLF5 on the SE of LINC00094.

Interestingly, verifying our recent findings on co-regulation between TCF3 and KLF5, silencing either one of these two TFs decreased the expression of the other (**Figure 2C**). Moreover, we further comprehensively analyzed H3K27ac and TCF3/KLF5 ChIP-seq data. Importantly, in the genome locus of TCF3, we identified multiple TCF3 and KLF5-binding peaks in a SE for TCF3 (**Figure 3A**). In the case of KLF5 locus, we also noted several TCF3 and KLF5 peaks at KLF5 SE (**Figure 3B**). Furthermore, the SE regions of these two TFs were highly specific to ESCC tumor samples and cell lines, as their signals were substantially weaker in normal samples (**Figure 3A-B**). To validate the ChIP-seq results, ChIP-qPCR was performed and the localization of TCF3 and KLF5 was quantified, and their enrichment was confirmed at their own and each other’s SEs (E1, E2, and E3) (Figure 3C-D). THZ1 and JQ1 inhibited this enrichment (**Figure 3C-D**). We next selected three candidate enhancer elements (E1, E2, and E3) each from TCF3 and KLF5 for luciferase reporter assays. Furthermore, silencing of either TCF3 or KLF5 decreased the reporter activity of the six enhancers, and overexpression of TCF3 or KLF5 increased significantly the reporter activity of the six enhancers (**Figure 3E-F**). The remaining cells from the luciferase reporter assays were collected for qRT-PCR, it was found that the mRNA levels of TCF3 and KLF5 decreased with the knockdown of the other and increased with the overexpression of the other (**Figure 3G-H**). Collectively, these results demonstrated that TCF3 and KLF5 co-exist in a CRC and regulate LINC00094 expression in a positive feedback loop.

**Figure 3.**
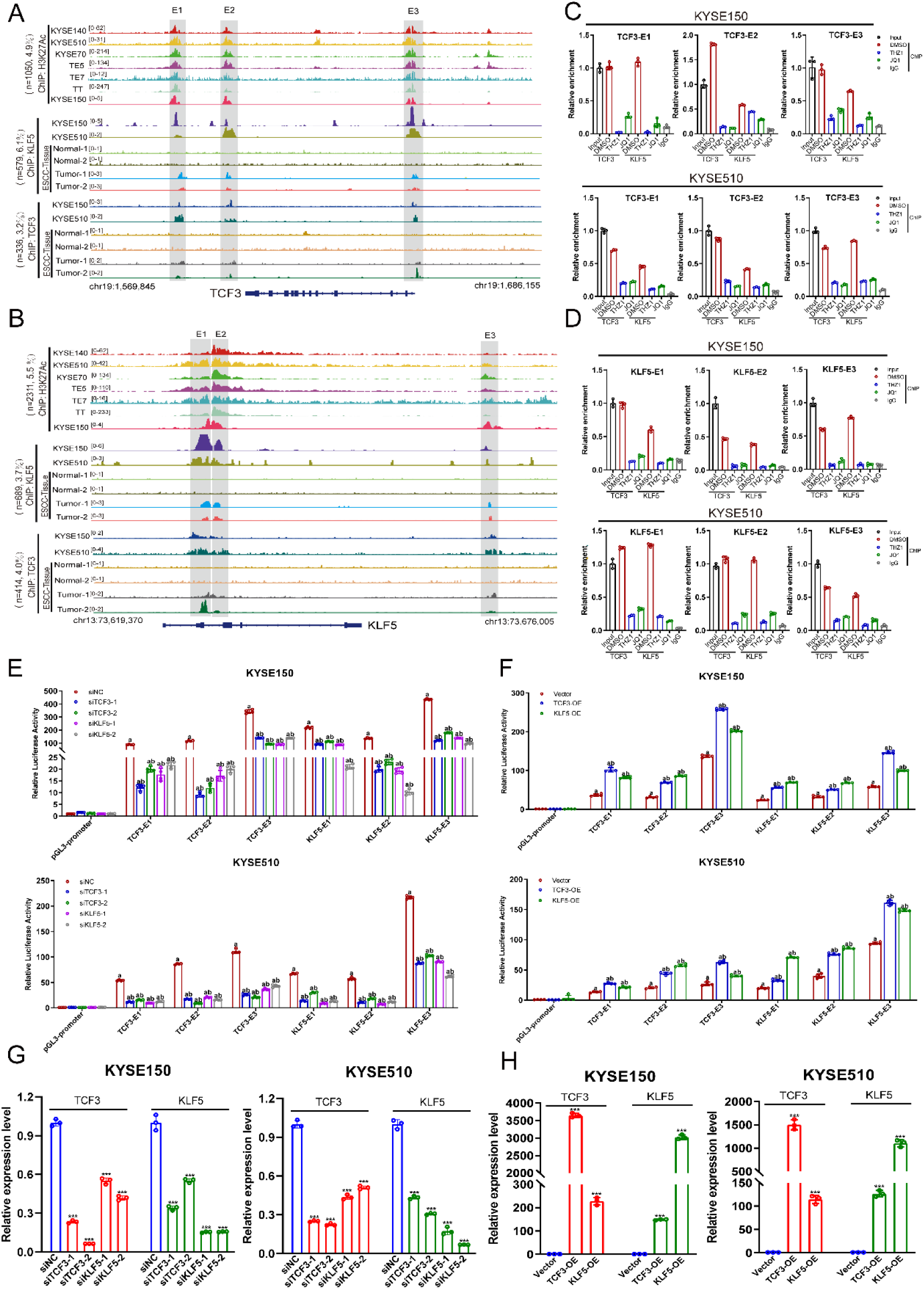
TCF3 and KLF5 regulate LINC00094 in a positive feedback loop. **A-B.** Localization profiles of TCF3, KLF5, and H3K27ac at the TCF3 (**A**) and KLF5 SE regions (**B**) in various types of ESCC cells and tissues. Gray shadings indicate the co-localization of TCF3, KLF5, and H3K27ac (n=the number of reads within the super-enhancer peaks of TCF3 or KLF5; percentage representation the number of reads within the super-enhancer peaks of the target molecules compared with the total number of reads in the ESCC cell lines or tumor samples). **C-D.** ChIP-qPCR experiments measuring TCF3 and KLF5 binding on the TCF3 (**C**) and KLF5 (**D**) SE segments (divided into enhancer 1, E1; enhancer 2, E2 and enhancer 3, E3) upon treatment with THZ1 (100 nM,12 h) and JQ1 (100 nM, 24 h). The technical triplicates in a representative experiment are shown and performed twice. Error bars indicate the mean ± SD from three replicates per group. IgG represents the NC antibody. **E.** TCF3 and KLF5 enhancer plasmids were transfected in KYSE150 and KYSE510 cells after knocking down TCF3 or KLF5. The cells were harvested 48 h later, and the reporter gene activity was measured. The firefly luciferase activity was normalized to Renilla luciferase activity, and the relative value from the cells transfected with the pGL3-promoter was set to 1. **F.** As described in **E**, the luciferase reporter gene assay after overexpression of TCF3 and KLF5 (TCF3-OE: TCF3 overexpression; KLF5-OE: KLF5 overexpression). Each value represents the mean ± SD, n≥3. a: siRNA or overexpression group compared with siNC or vector group, *P* < 0.01; b: pGL3-promoter-enhancer compared with pGL3-promoter-vector, *P* < 0.01. *P*-values were determined using a two-sided t-test. **G-H.** TCF3 and KLF5 expression in KYSE150 and KYSE510 cells detected by qRT-PCR after either TCF3 and KLF5 knockdown (**G**) or TCF3 and KLF5 overexpression (**H**). Each value represents the mean ± SD, n≥3. * *P* < 0.05, ** *P* < 0.01, *** *P* < 0.001. *P*-values were determined using a two-sided t-test.

### LINC00094, TCF3, and KLF5 co-regulate the transcription of each other

The above results showed that LINC00094 regulated the expression of TCF3 and KLF5 (**Figure 2**); meanwhile, TCF3 and KLF5 regulated the expression of LINC00094 in a positive feedback loop through SEs (**Figure 3** and Supplementary Figure 4). We speculated that LINC00094, TCF3, and KLF5 could form a new CRC. Specifically, we investigated RNA-seq data of ESCC cell lines from the Cancer Cell Line Encyclopedia (https://sites.broadinstitute.org/ccle) and noted that the expressions of TCF3, KLF5, and LINC00094 were prominently positively correlated (**Figure 4A**; Pearson correlation coefficients > 0.3). In addition, correlation analysis of RNA-seq expression profiles from 18 paired ESCC patients (PRJNA665149) also showed a positive correlation between the expression of LINC00094, TCF3 and KLF5 (Supplementary Figure 6A-B; Pearson correlation coefficients > 0.3). RNA-Seq dataset following LINC00094, TCF3, or KLF5 knockdown in KYSE150 and KYSE510 cells also identified the strong positive correlations between the expression levels of TCF3, KLF5, and LINC00094 (**Figure 4B**; Pearson correlation coefficients > 0.5). Importantly, RNA-seq data also showed that the knockdown of any single one of these three candidates decreased the expression of the other two (Supplementary Figure 6C). Subsequently, qRT-PCR was performed and the mRNA level silencing was quantified via siRNAs or shRNAs in ESCC cells to validate the RNA-seq results (**Figure 4C**). Consistently, the establishment of this CRC was further confirmed at the protein level using additional individual siRNAs or shRNAs (**Figure 4D**).

**Figure 4.**
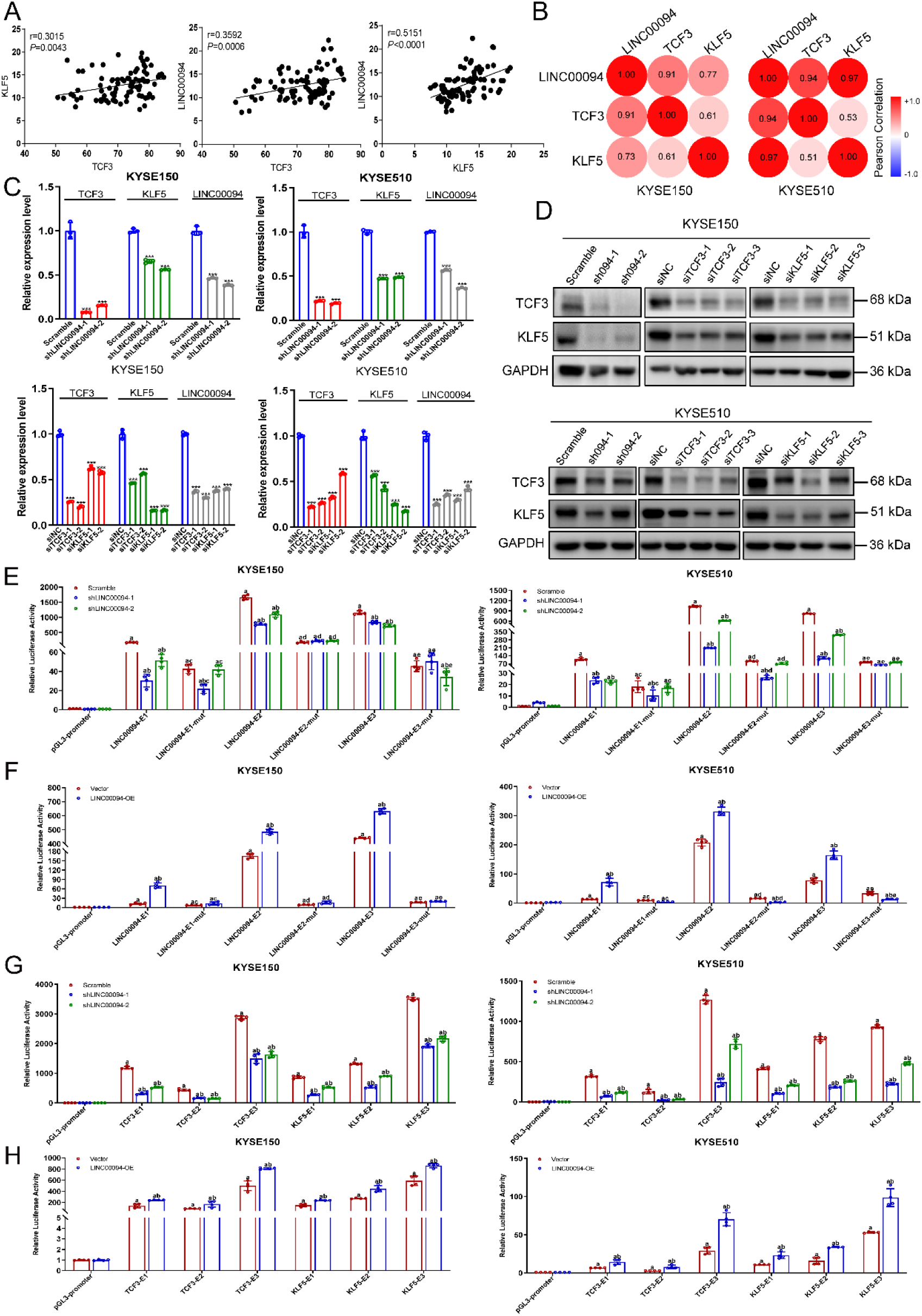
LINC00094, TCF3, and KLF5 regulate each other’s expression through SEs to form a CRC. **A.** Expression of LINC00094, TCF3, and KLF5 in ESCC cell lines (r>0 positive correlation, r<0 negative correlation). **B.** Heatmap of Pearson correlation coefficient among LINC00094, TCF3, and KLF5 expression levels in KYSE150 and KYSE510; RNA-seq data (red represents positive correlation, blue represents negative correlation). **C.** TCF3 or KLF5 expression in KYSE150 and KYSE510 cells detected via qRT-PCR after LINC00094 knockdown. Each value represents the mean ± SD, n≥3. * *P* < 0.05, ** *P* < 0.01, *** *P* < 0.001. **D.** Western blotting was performed to validate the co-regulation among LINC00094, TCF3, and KLF5 in KYSE150 and KYSE510 cells. The numbers denote the densitometric quantitation of band intensity, normalized to GAPDH levels. **E.** The LINC00094 stable knockdown cells were transfected with both the renilla plasmid and pGL3-promoter plasmid containing either wild-type or mutant LINC00094 E1/E2/E3 enhancers for 48 hours. The relative transcriptional reporter activity was analyzed by dual luciferase reporter gene assay. **F.** KYSE150 and KYSE510 cells were transfected with either empty vector or LINC00094 (LINC00094-OE) for 24 hours, and then co-transfected with both the renilla plasmid and pGL3-promoter plasmid containing either wild-type or LINC00094 E1/E2/E3 enhancers for another 48 hours. **G.** Relative luciferase activity of TCF3 and KLF5 enhancer reporter in LINC00094 stable knockdown cells. **H.** As described in **G**, the luciferase reporter activity assay after overexpression of LINC00094. Each value represents the mean ± SD, n≥3. a: siRNA or overexpression group compared with siNC or vector group, *P* < 0.01; b: pGL3-promoter-enhancer compared with pGL3-promoter-vector, *P* < 0.01; c: E1-mutant compared with E1, *P* < 0.05; d: E2-mutant compared with E2, *P* < 0.05; e: E3-mutant compared with E3, *P* < 0.05. *P*-values were determined using a two-sided t-test.

Whether LINC00094 can regulate its own as well as TCF3 and KLF5 expression through SEs is the key evidence for the involvement of LINC00094 in the formation of CRC. To confirm this, we first selected three enhancer elements (E1, E2, and E3) of LINC00094 SE for luciferase reporter assays. Robust reporter activities of E1, E2, and E3 were detected, and they were reduced upon silencing of LINC00094 (**Figure 4E**, Supplementary Figure 7A). The opposite result is obtained when LINC00094 is overexpressed (**Figure 4F, Supplementary Figure 7B**). We further performed site-directed mutagenesis to mutate TCF3 and KLF5-binding motif in E1, E2, and E3 elements. We found that knockdown of LINC00094 consistently decreased the reporter activity of both wild-type E1/E2/E3 elements, but failed to affect mutant E1/E2/E3 (**Figure 4E, Supplementary Figure 7A**). Similarly, overexpression of LINC00094 produced no detectable effect on the mutant enhancers (**Figure 4F, Supplementary Figure 7B**), confirming the direct regulation of LINC00094 on its own SE. Subsequently, three consecutive enhancers of TCF3 or KLF5 SE regions (**Figure 3A-B**) were individually cloned into pGL3-Promoter, all the enhancers reporter activities were strongly reduced upon LINC00094 silencing (**Figure 4G** and Supplementary Figure 7C), and LINC00094 overexpression significantly increased the reporter activity of the six enhancers (**Figure 4H** and Supplementary Figure 7D). These results suggested that LINC00094 regulated the expression of TCF3 and KLF5 by affecting their SE activity.

### LINC00094 recruits both TCF3 and KLF5 to form a new core regulation circuitry to regulate the expression of lipid metabolism-related genes

The above results suggested that LINC00094, TCF3, and KLF5 form a new CRC, but the mechanism by which LINC00094 is involved in this CRC is unknown. To further investigate the DNA binding of TCF3, KLF5, and LINC00094 in the nucleus, we performed a chromosome spreading assay. Interestingly, TCF3, KLF5, and LINC00094 were co-localized on the chromatin (**Figure 5A**). The localization of LINC00094 on the chromatin was reduced after knocking down TCF3 or KLF5 (**Figure 5A-B**). Sequence motif analysis confirmed that a large number of TCF3 and KLF5 motifs co-occupied the constitutive enhancers of LINC00094 SE (Supplementary Figure 4A). Moreover, a direct interaction between TCF3 and KLF5 in KYSE150 and KYSE510 cells was verified using co-IP assay (**Figure 5C**). Notably, re-ChIP experiments also showed that TCF3 was associated with KLF5 on the LINC00094 SE-binding sites (**Figure 5D**). These data characterized that TCF3 and KLF5 formed a complex that directly localizes at the SEs of LINC00094, thereby activating its transcription in ESCC cells. Importantly, we performed the RIP assay to pull down a TCF3-or KLF5-containing complex using either TCF3 or KLF5 antibody in ESCC cells, and qRT-PCR results showed that LINC00094 was strongly associated with TCF3 and KLF5, but not with the IgG antibody (**Figure 5E**). Furthermore, RNA-pulldown assay was conducted to complement the RIP assay. Western blotting results showed that a clear band corresponding to TCF3 or KLF5 was observed in both the input and LINC00094 fraction, whereas no band was observed in the LINC00094 antisense fraction (**Figure 5F**). Besides, downregulation of LINC00094 significantly reduced the binding between TCF3 and KLF5 (**Figure 5G**). The interaction between TCF3 and KLF5 no longer occurred after LINC00094 was degraded by RNase A (**Figure 5H**), suggesting that the interaction of TCF3 and KLF5 is directly dependent on LINC00094. These data demonstrated that LINC00094 participated in transcriptional regulation in ESCC by recruiting TCF3 and KLF5 to form a ternary complex, which forms a new CRC with TCF3 and KLF5.

**Figure 5.**
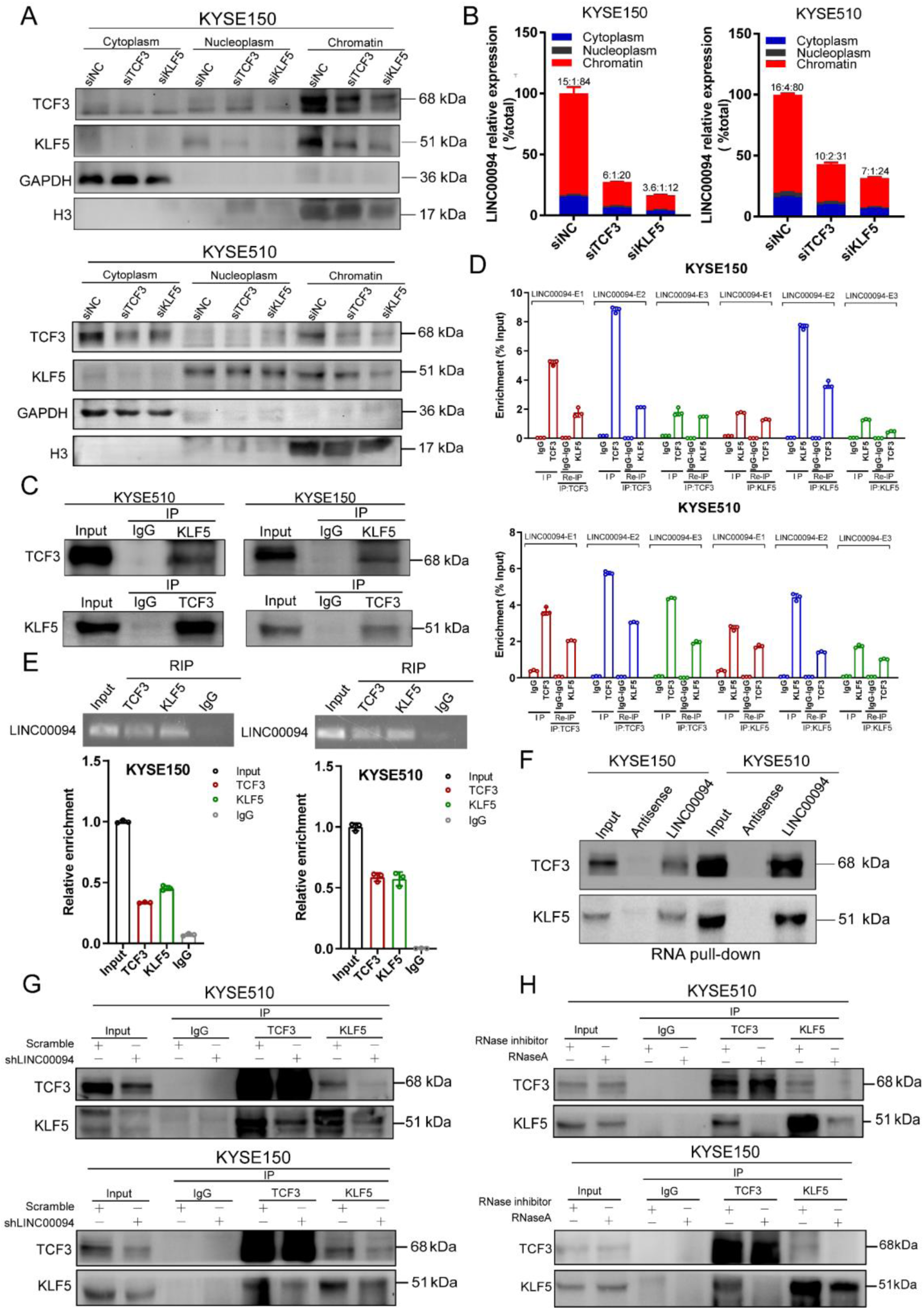
LINC00094 recruits both TCF3 and KLF5 to form a complex. **A.** Cytoplasm, nucleoplasm, and chromatin fractions from KYSE150 and KYSE510 cells were used for immunoblotting to detect TCF3 and KLF5. GAPDH and H3 were used as cytoplasmic and chromatin loading controls. **B.** The expression of LINC00094 in the cytoplasm, nucleoplasm, and chromatin fractions was detected via qRT-PCR after knocking down TCF3 and KLF5. LINC00094 localization was characterized by calculating the signal ratio of cytoplasm:nucleoplasm:chromatin. **C.** Immunoprecipitation of TCF3 or KLF5 using antibodies against endogenous TCF3 or KLF5 in KYSE150 and KYSE510 cells. **D.** Re-ChIP analysis of the localization of TCF3 and KLF5 on the indicated LINC00094 SE in KYSE150 and KYSE510 cells. **E.** RIP followed by RT-PCR analysis was used to detect the interaction between TCF3/KLF5 and LINC00094 in KYSE150 and KYSE510 cells. **F.** RNA-pulldown assay of biotin-labeled full-length LINC00094 RNA in KYSE150 and KYSE510 cells. Western blotting was used to detect TCF3 and KLF5 in the LINC00094 complex. Biotin-labeled LINC00094 antisense was used as the NC. **G.** Immunoprecipitation and western blotting were performed in stable LINC00094-knockdown KYSE150 and KYSE510 cells to detect the interaction of TCF3 and KLF5. **H.** Immunoprecipitation and western blotting was performed in RNaseA-treated KYSE150 and KYSE510 cells to detect the interaction of TCF3 and KLF5. Each value represents the mean ± SD, n≥3.

To validate this new CRC downstream targets, we first intersected RNA-seq data upon silencing LINC00094, TCF3, and KLF5 in ESCC cells. We identified a total of 464 common DEGs in KYSE150 cells and 301 common DEGs in KYSE510 cells (*P* <0.05, FC>1.2), most of them were SE-related genes (Supplementary Figure 8A). GO enrichment analysis revealed that common DEGs were enriched for processes involved in lipid metabolism, including lipid biosynthetic, phospholipid metabolic, and regulation of lipid metabolic process (**Figure 6A** and Supplementary Figure 8B). Moreover, depletion of LINC00094/TCF3/KLF5 consistently reduced total lipid droplet levels, especially triple knockdown of the three molecules, suggesting decreased lipid storage (**Figure 6B-D** and Supplementary Figure 9). Meanwhile, ectopic expression of LINC00094 reversed the reduction of lipid droplet content caused by LINC00094/TCF3/KLF5-depletion (**Figure 6B-D** and Supplementary Figure 9). Subsequently, to understand whether the CRC affected the phenotype of ESCC by regulating fatty acid synthesis, we further analyzed the RNA-seq data from our study (LINC00094, TCF3, or KLF5 knockdown). We identified some lipid metabolism-related genes that were significantly differentially expressed, most of which were fatty acid synthesis-related enzymes and TFs, such as ACSS2, PDK2, PPARD, and CEBPA (Supplementary Figure 10A). Notably, qRT-PCR was performed, and the mRNA level upon LINC00094, TCF3, and KLF5 silencing via siRNAs or shRNAs in KYSE150 and KYSE510 were quantified to validate the RNA-seq results. The results showed that the expression of most of the selected lipid metabolism-related genes were highly correlated with the expression of TCF3, KLF5, and LINC00094 (**Figure 6E-F** and Supplementary Figure 10B-C). In particular, when TCF3, KLF5, and LINC00094 were silenced together, the expression of lipid metabolism-related genes changed most significantly (**Figure 6E-F** and Supplementary Figure 10B-C).

**Figure 6.**
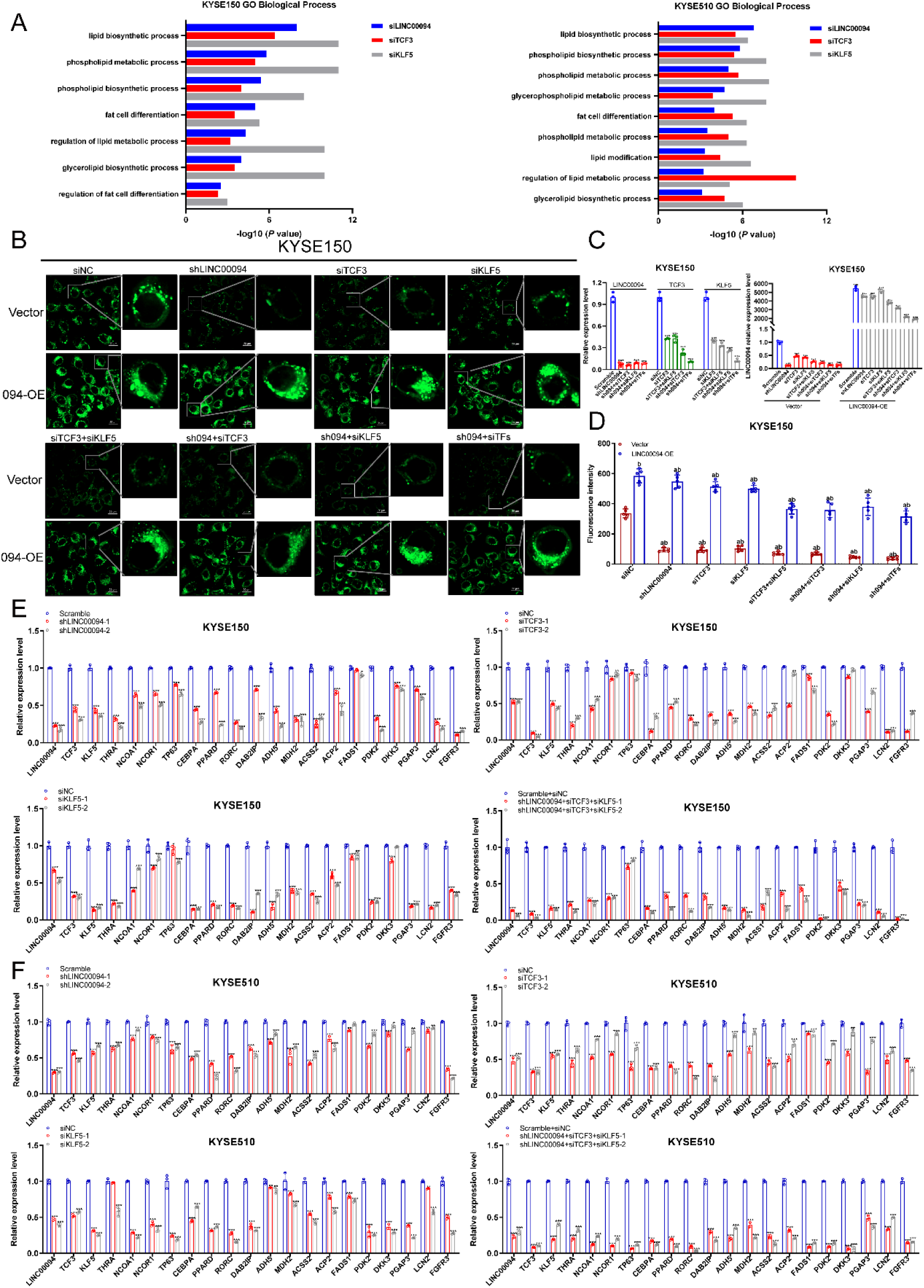
LINC00094, TCF3, and KLF5 cooperate to regulate the lipid metabolism pathway in ESCC. **A.** GO enrichment analysis of common differentially downregulated genes upon LINC00094, TCF3, and KLF5 silencing in KYSE150 and KYSE510 cells. **B.** Confocal images of staining of lipid droplets after either LINC00094/TCF3/KLF5 knockdown or combined with full-length LINC00094 overexpression in KYSE150 cells (sh094: shLINC00094; siTFs: siTCF3+siKLF5; 094-OE: LINC00094 overexpression). Scale bar, 20 μm. **C.** qRT-PCR analysis was performed to detect the mRNA levels of LINC00094, TCF3, and KLF5 after knockdown or overexpression in different treatment groups of lipid droplet staining assay. Each value represents the mean ± SD, n≥3. * *P* < 0.05, ** *P* < 0.01, *** *P* < 0.001. **D.** Quantitative analysis of lipid droplet staining based on the confocal images; Mean ± SD are shown, n = 5, as the number of microscopic vision. a: siRNA or overexpression group compared with siNC or vector group, *P* < 0.01; b: LINC00094-OE compared with vector, *P* < 0.01. **E-F.** qRT-PCR analysis was performed to detect the low differential expression of lipid metabolism-related genes after knocking down LINC00094, TCF3, and KLF5 in KYSE150 (**E**) and KYSE510 (**F**) cells. Each value represents the mean ± SD, n≥3. * *P* < 0.05, ** *P* < 0.01, *** *P* < 0.001. *P*-values were determined using a two-sided t-test.

### LINC00094, TCF3, and KLF5 cooperate to activate lipid metabolism related-genes’ expression through SEs in ESCC

Having established the molecular basis of the SE activation of LINC00094, as well as a new CRC formed by LINC00094/TCF3/KLF5 in ESCC cells, we next investigated the mechanisms underlying the functional impact of this CRC on fatty-acid metabolism identified earlier. Interestingly, we analyzed ChIP-seq data of H3K27ac, TCF3, and KLF5 in ESCC cell lines and tissues. Indeed, SE-associated genes involved in lipid metabolism were annotated (Supplementary Figure 10A). Several lipid metabolism-related genes regulated by SEs in seven ESCC cell lines (59%, 24/40), whereas 80% (32/40) of the lipid metabolism-related genes in KYSE150 and KYSE510 cells were SE-related genes (**Figure 7A**). We hypothesized that lipid metabolic processes in ESCC are closely related to the regulation of SEs. To confirm this hypothesis, we treated ESCC cells with THZ1 and JQ1 and found that lipid droplet synthesis was significantly reduced at low drug concentrations (50 nM, 12 h) (**Figure 7B-C** and Supplementary Figure 11). Further analysis of the above genes co-regulated by LINC00094/TCF3/KLF5 in combination with ChIP-seq data revealed a numbered of well-defined lipid metabolism-related genes had super-enhancers (e.g. ACSS2, PDK2, CEBPA, and PPARD) ^[34, 35, 40–44]^ in two ESCC cell lines (Figure 7D). Importantly, we further comprehensively analyzed H3K27ac and TCF3/KLF5 ChIP-seq data. In the case of ACSS2/PDK2/CEBPA/PPARD locus, we identified multiple TCF3 and KLF5-binding peaks in SEs for these molecules (**Figure 7E**). Subsequently, ChIP-qPCR confirmed the enrichment of TCF3 and KLF5 at SEs of ACSS2/PDK2/CEBPA/PPARD, THZ1 and JQ1 inhibited the interaction of TCF3 and KLF5 with these SEs (**Figure 7F** and **Supplementary Figure S12**). Additionally, qRT-PCR analysis after RNA extraction from xenograft mouse tumors showed that the expression of ACSS2/PDK2/CEBPA/PPARD was also significantly reduced in the LINC00094 knockdown group (Supplementary Figure 13). These results suggested that the new CRC composed of LINC00094, TCF3, and KLF5 directly regulates the expression of key lipid metabolism-related genes through SEs.

**Figure 7.**
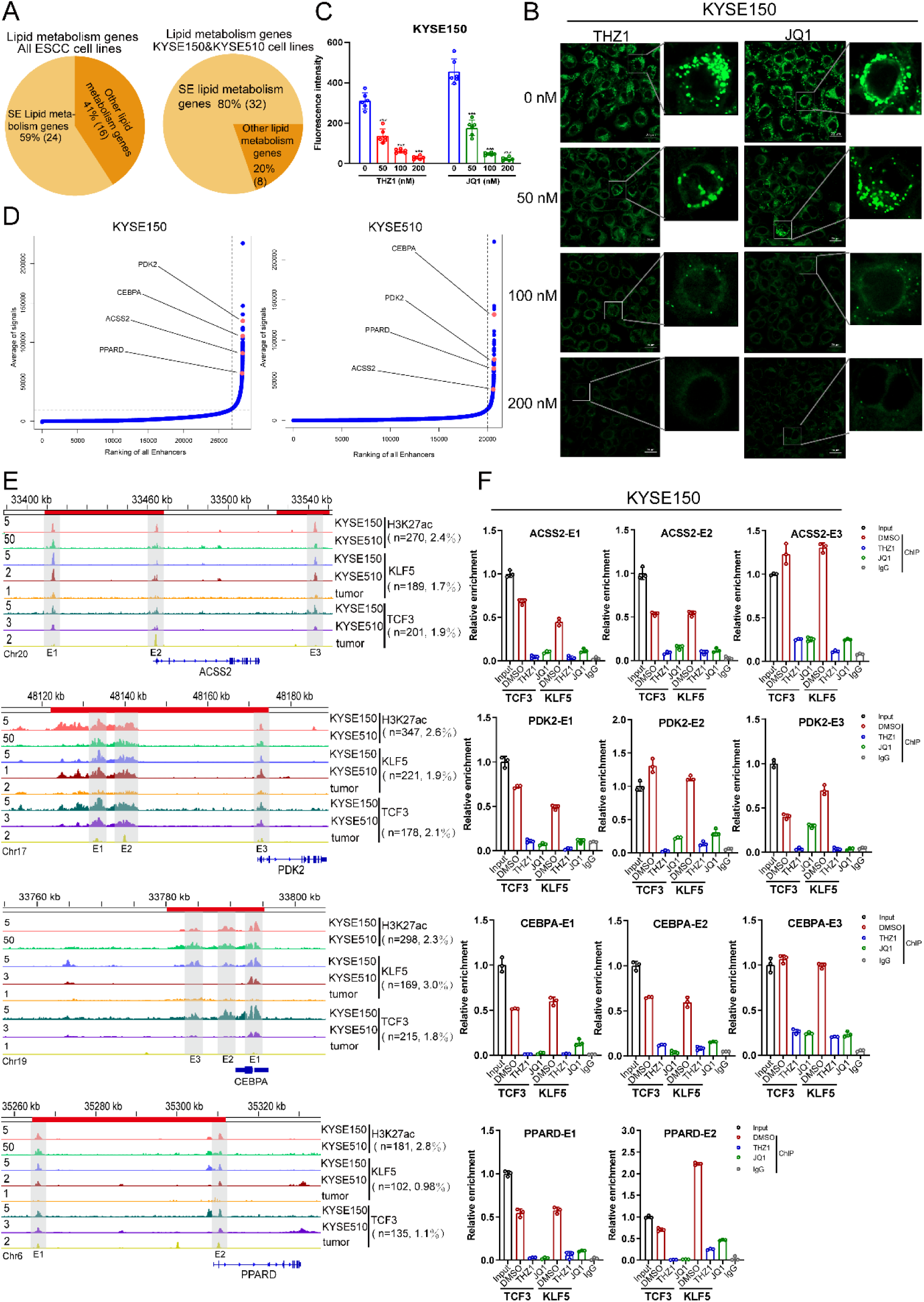
The novel CRC formed by LINC00094, TCF3, and KLF5 directly regulate lipid metabolism-related genes transcription through SE. **A.** The pie chart displays SE-associated lipid metabolism genes in ESCC. **B.** Confocal images of staining of lipid droplets after THZ1 and JQ1 treatment (0, 50, 100, 200 nM, 12 h) in KYSE150 cells. **C.** Quantitative analysis of lipid droplet staining based on the confocal images; Mean ± SD are shown, n = 6, as the number of microscopic vision. *** *P* < 0.001. *P*-values were determined using a two-sided t-test. **D.** Hockey stick plots of KYSE150 and KYSE510 showing normalized, rank-ordered H3K27ac input signals for SE-associated genes. **E.** IGV showing normalized ChIP-seq signals for the H3K27ac, TCF3, and KLF5 at the ACSS2/PDK2/CEBPA/PPARD locus in ESCC cells and tissues. Upper red bars denote SE regions. Gray shadings indicate the co-occupancy of TCF3, KLF5, and H3K27ac at constituent enhancers within SEs (n=the number of reads within the super-enhancer peaks of ACSS2, PDK2, CEBPA, or PPARD; percentage representation the number of reads within the super-enhancer peaks of the target molecules compared with the total number of reads in the ESCC cell lines or tumor samples). **F.** ChIP-qPCR experiments measuring TCF3 and KLF5 binding on the ACSS2/PDK2/CEBPA/PPARD SE segments upon treatment with THZ1 (100 nM,12 h) and JQ1 (100 nM, 24 h). The technical triplicates in a representative experiment are shown and performed twice. Error bars indicate the mean ± SD from three replicates per group. IgG represents the NC antibody.

To further explore the clinical significance of the new CRC, we used the LASSO Cox regression algorithm assay in the training group (GSE53625, *n* =179) to construct a signature for evaluating the prognosis of patients with ESCC. Finally, 28 DEGs were chosen to establish a prognostic signature, and the risk score was calculated (**Figure 8A-B**). The patients in the training group were classified into two groups, namely, the high-risk and low-risk groups using the median risk score as the demarcation value. Kaplan-Meier analysis showed that the low-risk group had a significantly longer survival time than the high-risk group (*P*< 0.001) (**Figure 8C**). The model established in this study was also meaningful in the validation set (TCGA, *n* = 84); the results showed significant differences in OS between high-and low-risk groups (*P*< 0.005) (**Figure 8D**). Moreover, among the 28 genes in this signature, 21 genes were regulated by LINC00094, TCF3, and KLF5 (Supplementary Figure 14A-B). GO analysis of these 21 genes was also enriched for lipid metabolism-related pathways, including lipid transport and binding (Supplementary Figure 14C). Univariate COX analysis showed that most of these genes were significant for survival (Supplementary Figure 14D). Collectively, these results highlighted that the CRC formed by LINC00094/TCF3/KLF5 regulates the expression of lipid metabolism-related genes through SEs, thereby affecting ESCC phenotypes.

**Figure 8.**
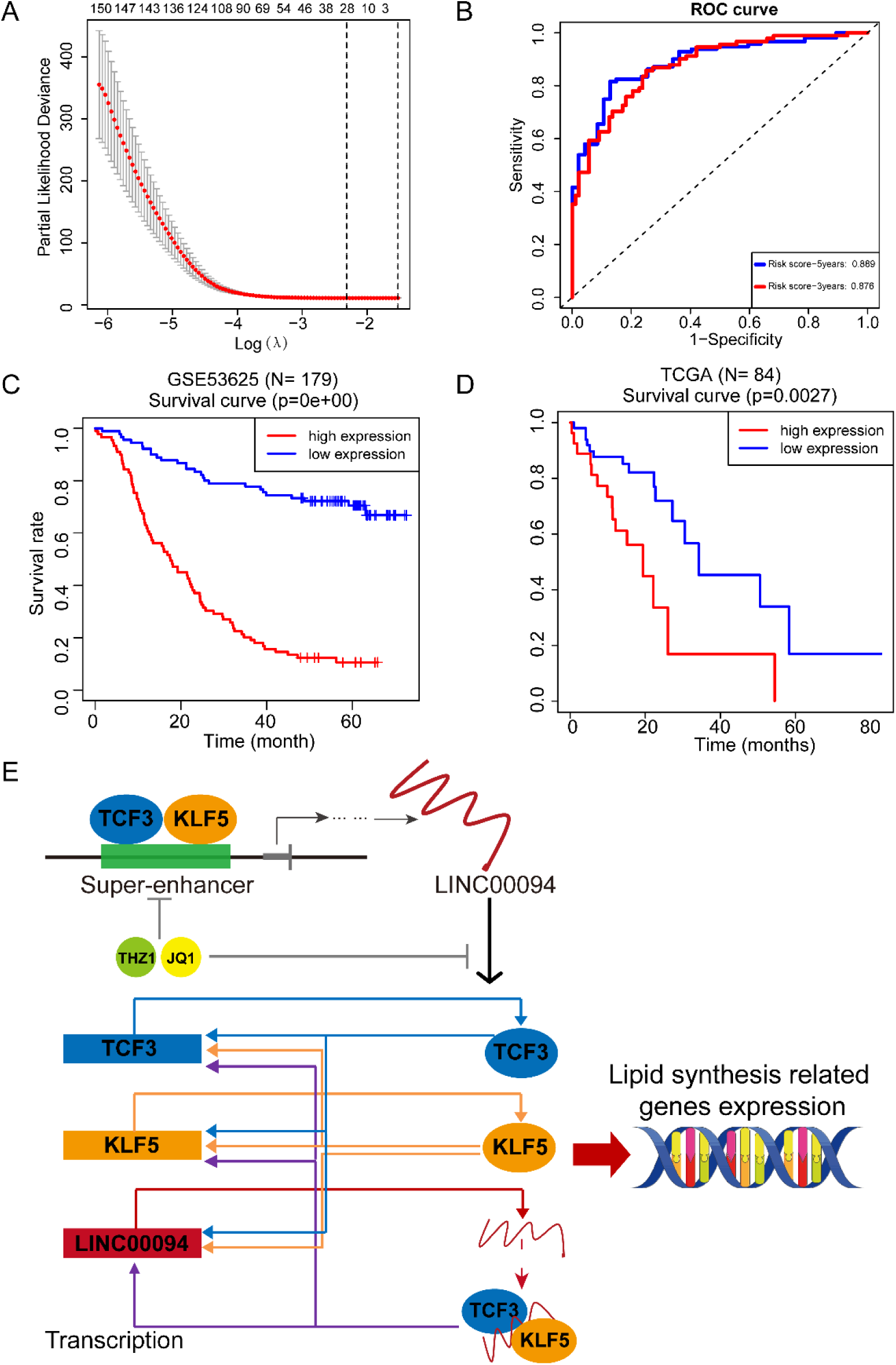
Identification of LINC00094-related signatures using LASSO regression algorithm in ESCC. **A.** The lines in different colors represent the trajectory of the correlation coefficient of different factors in the model with the increase of Log Lamda. **B.** The point with the smallest cross-verification error corresponds to the number of factors included in the LASSO regression model. **C-D.** Kaplan-Meier analysis of the prognostic risk model selected using LASSO Cox regression of patients with ESCC in the training (**C,** GSE53625, n=179) and independent validation cohorts (**D,** TCGA, n=84). **E.** The transcriptional regulatory model of LINC00094, TCF3, and KLF5 form a new CRC. TCF3 and KLF5 co-localize at LINC00094 SE regions to activate LINC00094 expression. LINC00094 in turn regulates the expression of TCF3 and KLF5 through recruits both TCF3 and KLF5 and forms a protein/RNA complex, which forms a CRC with TCF3 and KLF5 to activate each other’s and lipid metabolism-related genes transcription through SEs. This transcriptional regulation pattern can be effectively repressed by THZ1 and JQ1.

## Discussion

The regulatory mechanism of lncRNAs with SEs and CRC have not been studied extensively. The present study provides compelling evidence to highlight closely cooperative machinery among SEs, lncRNA, and master TFs that contribute to ESCC malignancy. We established that TCF3 and KLF5 transcriptionally activated LINC00094 expression through direct co-localization at its SE elements, and TCF3 and KLF5 co-existed in a CRC to promote LINC00094 expression in a positive feedback loop. LINC00094 also regulated the expression of TCF3 and KLF5 by recruiting both TCF3 and KLF5, forming a protein/RNA complex, which developing a new CRC with TCF3 and KLF5 to activate each other’s transcription. The complex regulations resulted in the high expression of LINC00094, TCF3, and KLF5, promoting ESCC tumor progression by activating transcriptional expression of lipid metabolism-related genes. The formation of this new CRC is dependent on LINC00094, and the SE inhibitors (THZ1 and JQ1) can effectively inhibit the formation of this CRC and its regulation of downstream genes (**Figure 8E**).

We first confirmed that LINC00094 played a transcriptional regulatory role by participating in the CRC. So far, there is no evidence to prove that lncRNAs can form a CRC. A key mechanism of lncRNAs in transcriptional regulation is to act as a scaffold to recruit other transcriptional regulators ^[45]^. LINC00094, which is located in the nucleus, promoted the expression of TCF3 and KLF5 by affecting their SE activity. LINC00094 could recruit TCF3 and KLF5 to form a ternary complex, and LINC00094 was necessary to form this complex. The expressions of LINC00094, TCF3, and KLF5 were prominently positively correlated. Our research suggested that LINC00094 was involved in TCF3/KLF5-mediated CRC by recruiting TCF3 and KLF5 to form a ternary complex. It should be noticed that the downstream targets of the CRC were also abundantly enriched in pathways related to RNA polymerase Ⅱ (RNA pol Ⅱ), such as RNA pol Ⅱ transcription factor/cofactor binding and RNA pol Ⅱ transcription factor complex (Supplementary Figure 8B). As expected, RIP and RNA-pulldown experiments proved that LINC00094 directly interacted with RNA pol Ⅱ (Supplementary Figure 15). This result indicated that the CRC might combine with RNA pol Ⅱ. Recent studies have shown that RNA polymerase II recruits many transcriptional regulatory factors at SEs and forms transcriptional regulatory machinery through phase separation to regulate downstream gene transcription efficiently ^[46–49]^. Phase separation is the basis for forming membrane-less organelles in cells ^[49, 50]^. Many biological macromolecules, such as proteins and nucleic acids can fuse and gather to form liquid-like structures that perform specific biological functions ^[50–52]^. Phase separation is driven by collective, weak, and multivalent interactions between biological macromolecules. Some special protein domains (such as SH3, PRM, and bHLH), intrinsically disordered regions, and RNA repetitive sequences (such as CUG, CAU, and GGGGCC) play an important role in the effect of the multivalent interaction ^[51, 53–56]^. LINC00094 is a non-coding RNA with 2992 nucleotides, containing a large number of repetitive sequences. At the same time, LINC00094 recruits master TFs and binds to RNA pol Ⅱ, which indicates that LINC00094 may form transcriptional regulatory machinery through phase separation in the SE region and efficiently regulate the transcription of downstream genes. This is a new mechanism for lncRNA to participate in transcriptional regulation.

Lipid metabolism, particularly the synthesis of fatty acids, is an essential cellular process that converts nutrients into metabolic intermediates for membrane biosynthesis, energy storage, and signaling molecule generation ^[57, 58]^. Reprogramming of the lipid metabolism is a newly recognized hallmark of malignancy ^[57, 59]^. Several decades ago, tumors were found to synthesize fatty acids de novo and exhibit a shift toward fatty acid synthesis ^[60, 61]^. Increased lipid uptake, storage, and de novo fatty acid synthesis are a result of various cancers and contribute to rapid tumor growth and proliferation ^[20, 57, 58, 62–65]^. However, it is unclear how fatty acid metabolism is dysregulated in ESCC. Our data demonstrated the cooperative regulation of LINC00094/TCF3/KLF5 on the transcription of metabolism-related genes associated with SEs. More importantly, most of the metabolism-related genes were fatty acid synthesis-related enzymes and TFs. For example, the expression of ACSS2, PDK2, PPARD, and CEBPA was significantly downregulated upon silencing LINC00094/TCF3/KLF5. Acetate is converted to acetyl-CoA by ACSS2, making acetate an important molecule for lipid synthesis and histone acetylation ^[40, 41, 57]^. Pyruvate dehydrogenase kinase 2 (PDK2) is an enzyme that plays a key role in glycolipid metabolism ^[42, 43]^. CEBPA and PPARD are key TFs that control the synthesis and differentiation of adipocytes ^[33, 66–69]^. It is worth noting that KLF5 is a key factor in regulating lipid synthesis ^[32, 33]^. We previously found that KLF5, SREBF1, and TP63 co-regulate lipid synthesis in squamous cell carcinoma ^[20]^. Many studies in recent years have shown that increased lipid synthesis is closely related to cisplatin resistance. Cisplatin is an important chemotherapeutic agent against various solid tumors, but resistance often limits its usage. Wen et al. found that high ACSS2 expression is observed in cisplatin-resistant bladder cancer patient tissues, and bladder cancer cells are more sensitive to cisplatin after inhibiting the expression of ACSS2 ^[70]^. Another study has shown that the accumulation of lipid droplets in colorectal cancer can lead to a cisplatin-resistant phenotype ^[71]^. Here, we showed that depletion of LINC00094/TCF3/KLF5 inhibits the expression of key enzymes and related TFs for fatty acid synthesis, thereby consistently reducing total lipid droplet levels. Therefore, the CRC composed of LINC00094, TCF3, and KLF5 targeting lipid metabolism is likely to be further utilized as a potential target for ESCC therapy. However, the deeper molecular mechanisms need to be further studied and explored.

## Conclusion

A new CRC containing non-coding RNA is identified in ESCC tissues and cells. The CRCs constituted LINC00094, TCF3, and KLF5, which regulate the transcription of downstream genes related to lipid metabolism in the SE region, promote lipid synthesis, and lead to malignant progression of ESCC. The development of CRC inhibitors, similar to JQ1 and THZ1, will have important clinical significance for the targeted therapy of patients with ESCC.

### Abbreviations

AUC: area under the curve
ChIP-seq: chromatin immunoprecipitation
co-IP: co-immunoprecipitation
CRC: core transcriptional regulation circuitry
DEG: differentially expressed gene
ESCC: esophageal squamous cell carcinoma
GEO: Gene Expression Omnibus
GO: gene ontology
H3K27ac: Acetylation at the 27th lysine residue of the histone H3 protein
KLF5: kruppel like factor 5
IGV: integrative genomics viewer
LASSO: least shrinkage and selection operator
lncRNA: long non-coding RNA
NC: Negative control
OS: overall survival
RIP: RNA binding protein immunoprecipitation
ROC: receiver operating characteristic curve
RT-qPCR: reverse transcriptase-quantitative real-time PCR
SE: super-enhancer
SEM: standard error of the mean
TCF3: transcription factor 3
TF: transcription factor

## Acknowledgments

The authors are indebted to all the donors whose names were not included in the author list, but who participated in our study.

## Author contributions

The study design: L.-Y. Xu, E.-M. Li, and C.-Q. Li; The in vitro and in vivo experiment and performed data analysis: L. Peng, J.-X. Chen, Y. Chen, Q.-Y. Wang, L.-D. Liao and L. Wan; The animal experiments: L. Peng and R.-Y. Li; The clinical data analysis: Y. Chen; Writing the manuscript: L. Peng, Q.-Y. Wang, C.-Q. Li, L.-Y. Xu and E.-M. Li. All the authors read and approved the final manuscript.

## Availability of data and materials

Our RNA-seq and ChIP-seq data in this study have been deposited in NCBI’s Sequence Read Archive under accession numbers PRJNA791409 and PRJNA792889.

## Ethics approval

Ethical consent was approved by the Committees for Ethical Review of Research involving Human Subjects at Shantou University Medical College, approval number is SUMC-2021-68. Written informed consent was obtained from each patient before sample collection. The animal experiments were approved by the Use Committee for Animal Care at Shantou University Medical College, approval number is SUMC-2021338.

## Consent for publication

Not applicable.

## Supplementary figures

**Figure S1.**
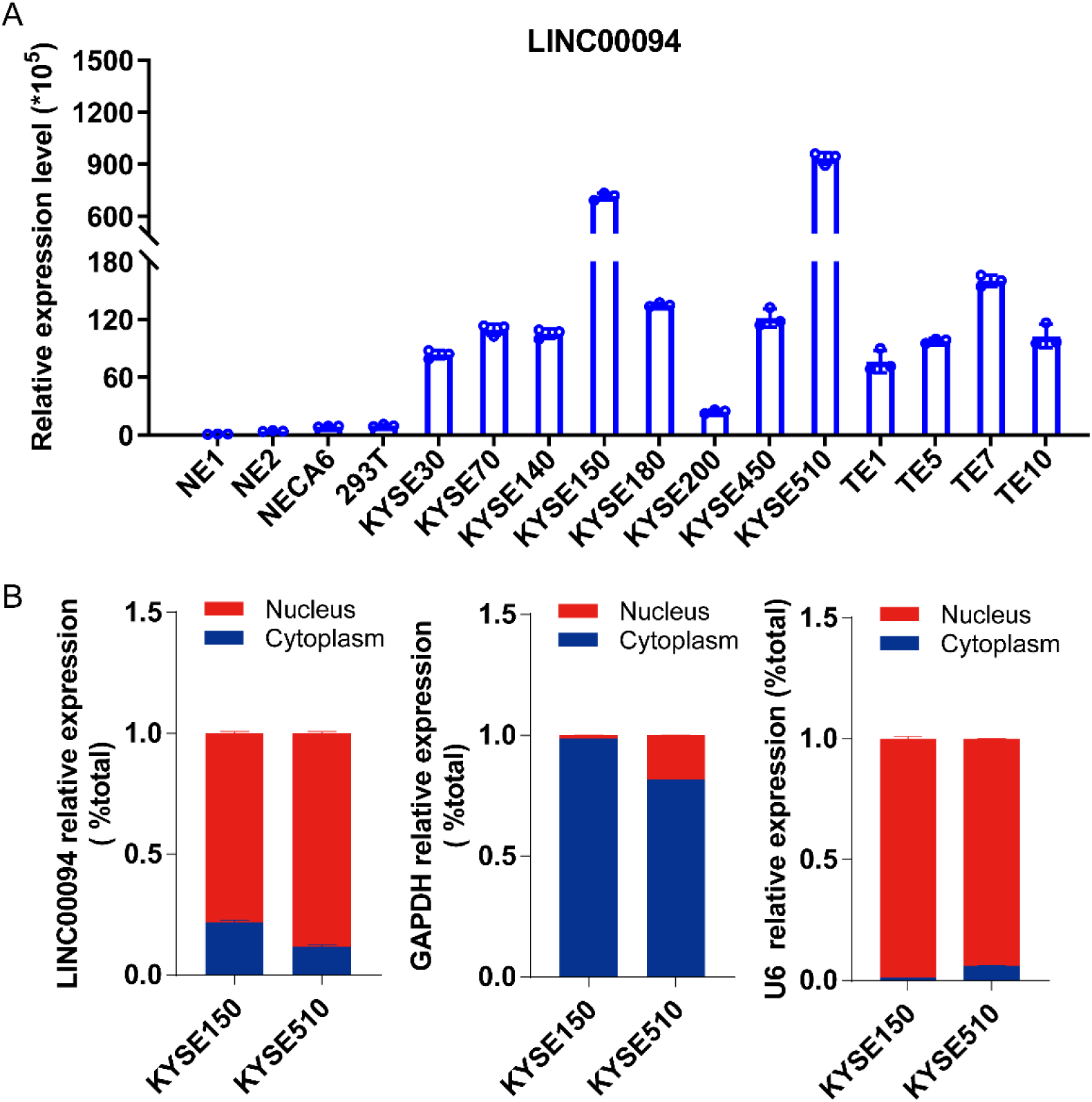
LINC00094 is mainly localized in the nucleus. **A.** Relative expression of LINC00094 in normal esophageal epithelial cells and ESCC cells. **B.** Relative expression of LINC00094 in nuclear and cytoplasmic RNA samples was detected via qRT-PCR in KYSE150 and KYSE510 cells. GAPDH and U6 were used as the cytoplasmic and nuclear loading controls, respectively.

**Figure S2.**
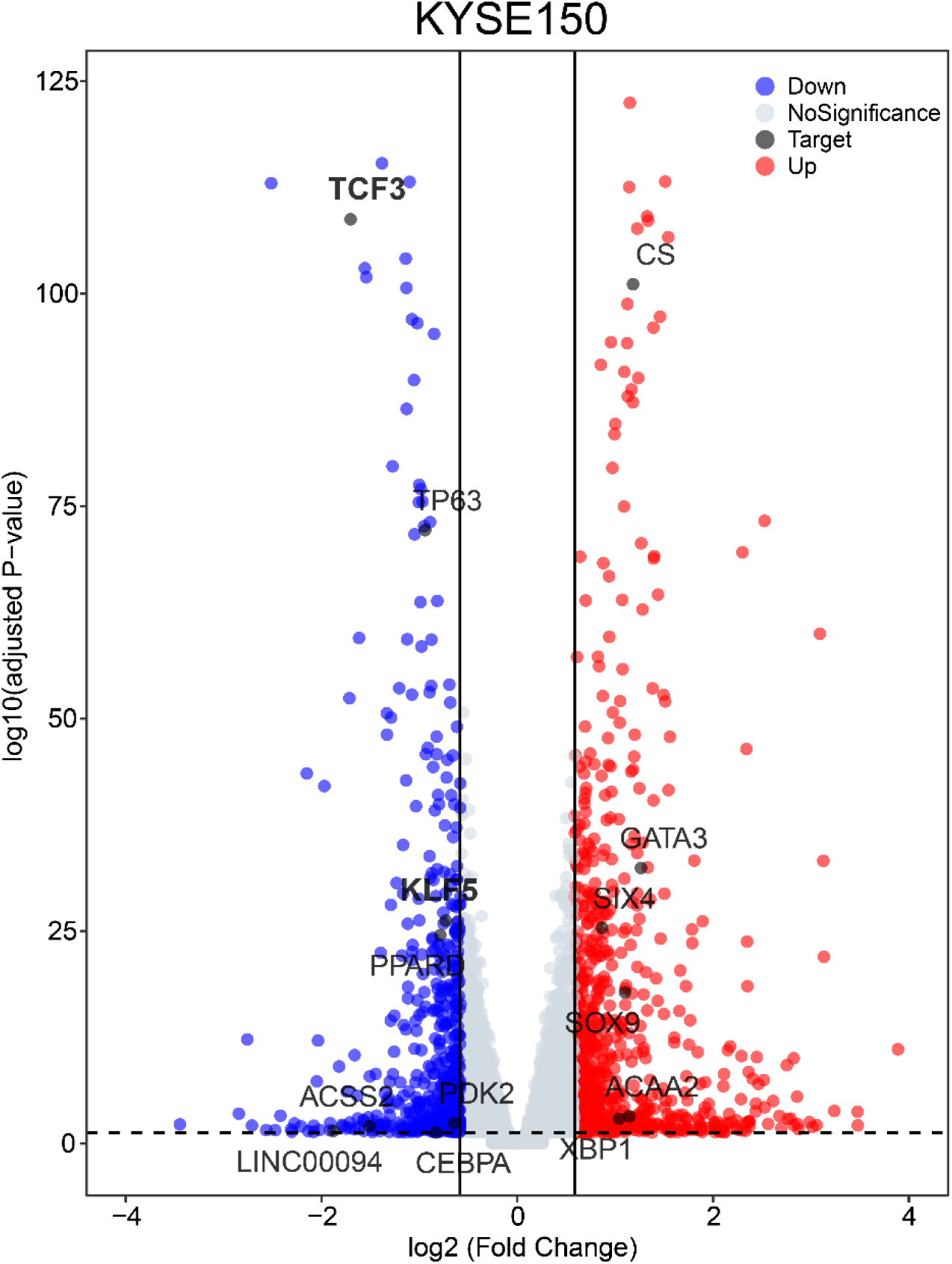
The volcano plot of DEGs in KYSE150 cells after LINC00094 knockdown. Related to Figure 1. Red represents the differentially upregulated genes and blue represents the differentially downregulated genes.

**Figure S3.**
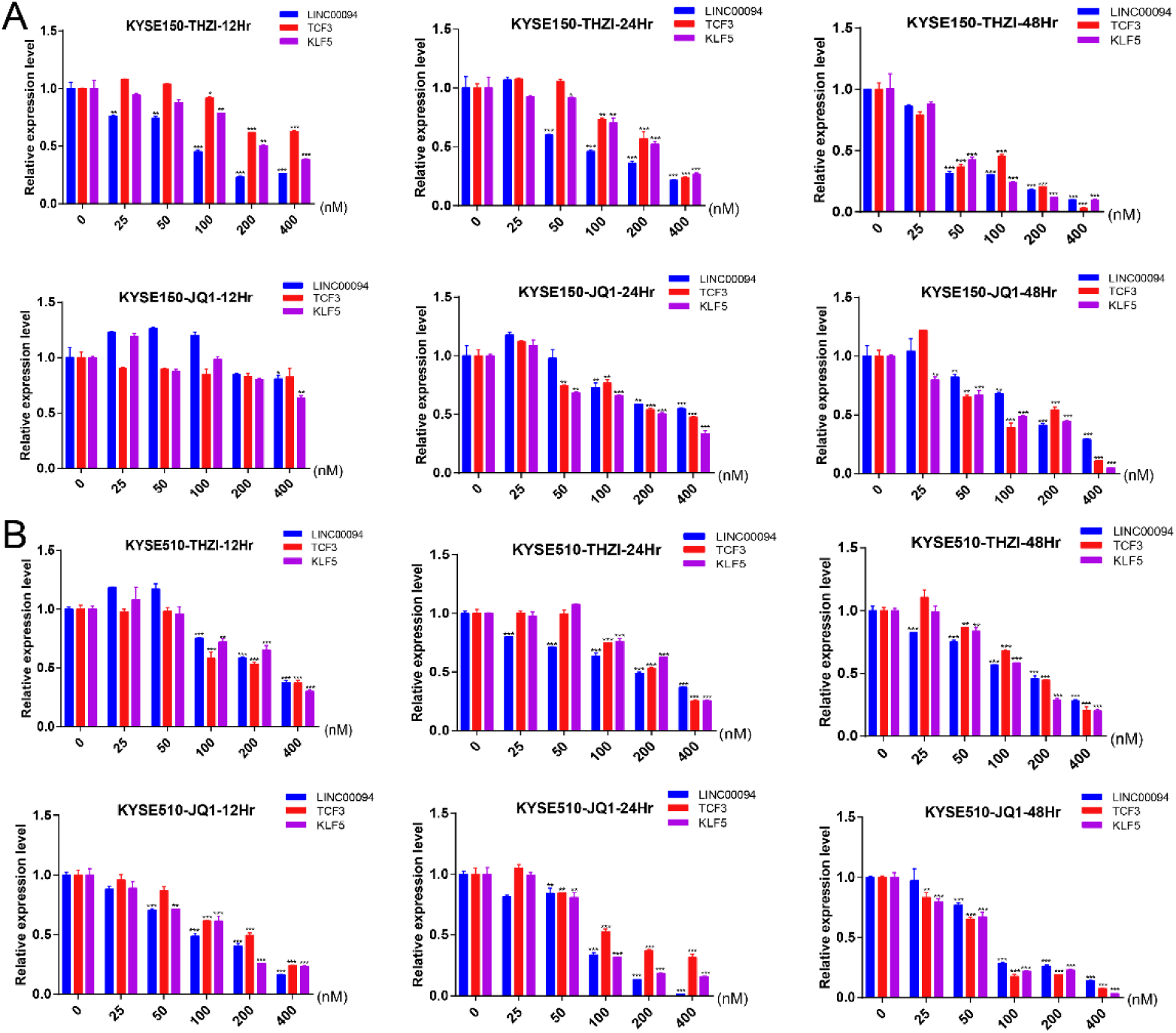
qRT-PCR of LINC00094, TCF3, and KLF5 after SE inhibitor treatment. KYSE150 **(A)** and KYSE510 **(B)** cells were treated with THZ1 and JQ1 to detect mRNA expression levels of LINC00094, TCF3, and KLF5. Each value represents the mean ± SD. The data are representative of at least two independent experiments. * *P* < 0.05, ** *P* < 0.01, *** *P* < 0.001.

**Figure S4.**
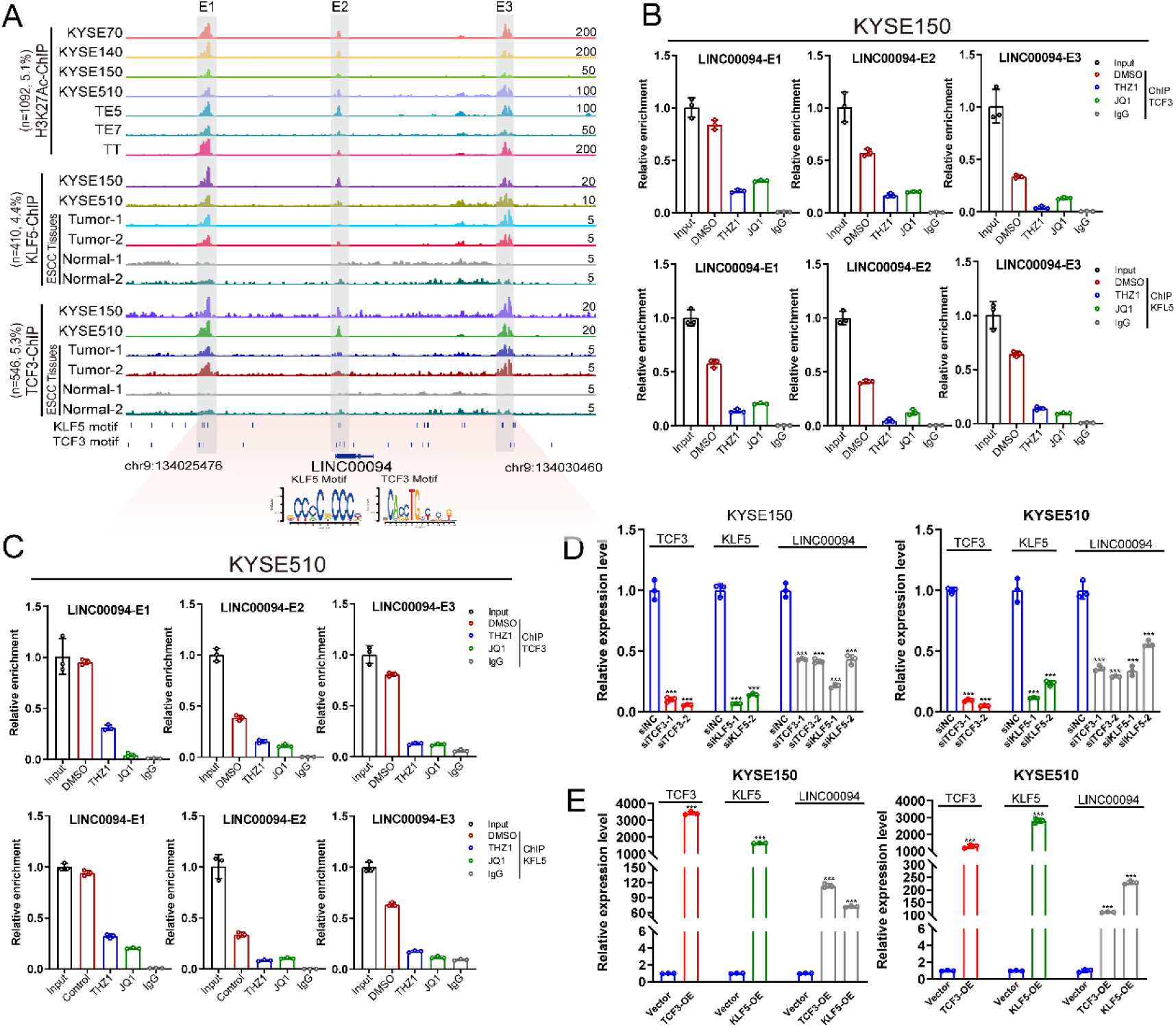
TCF3 and KLF5 directly regulate LINC00094 transcription through SE. **A.** Occupancy profiles of TCF3, KLF5, and H3K27ac at the LINC00094 SE regions in various types of ESCC cells and tissues. Gray shadings indicate the co-occupancy of TCF3, KLF5, and H3K27ac and three constituent enhancers (E1, E2, and E3) within the SE. TCF3 and KLF5 motifs occupy E1, E2, and E3 enhancer loci (n=the number of reads within the super-enhancer peaks of LINC00094; percentage representation the number of reads within the super-enhancer peaks of the target molecules compared with the total number of reads in the ESCC cell lines or tumor samples). **B-C.** ChIP-qPCR analysis in KYSE150 (**B**) and KYSE510 (**C**) for the enrichment of TCF3 and KLF5 at the LINC00094 SEs (divided into enhancer 1, E1; enhancer 2, E2; enhancer 3, E3). The means of technical triplicates are shown in a representative experiment, performed twice. Error bars indicate mean ± SD from three replicates per group. IgG represents the NC antibody. **D-E.** LINC00094, TCF3, and KLF5 expression in KYSE150 and KYSE510 cells detected via qRT-PCR after either TCF3 and KLF5 knockdown (**D**) or TCF3 and KLF5 overexpression (**E**). Each value represents the mean ± SD, n≥3. * *P* < 0.05, ** *P* < 0.01, *** *P* < 0.001. P-values were determined using a two-sided t-test.

**Figure S5.**
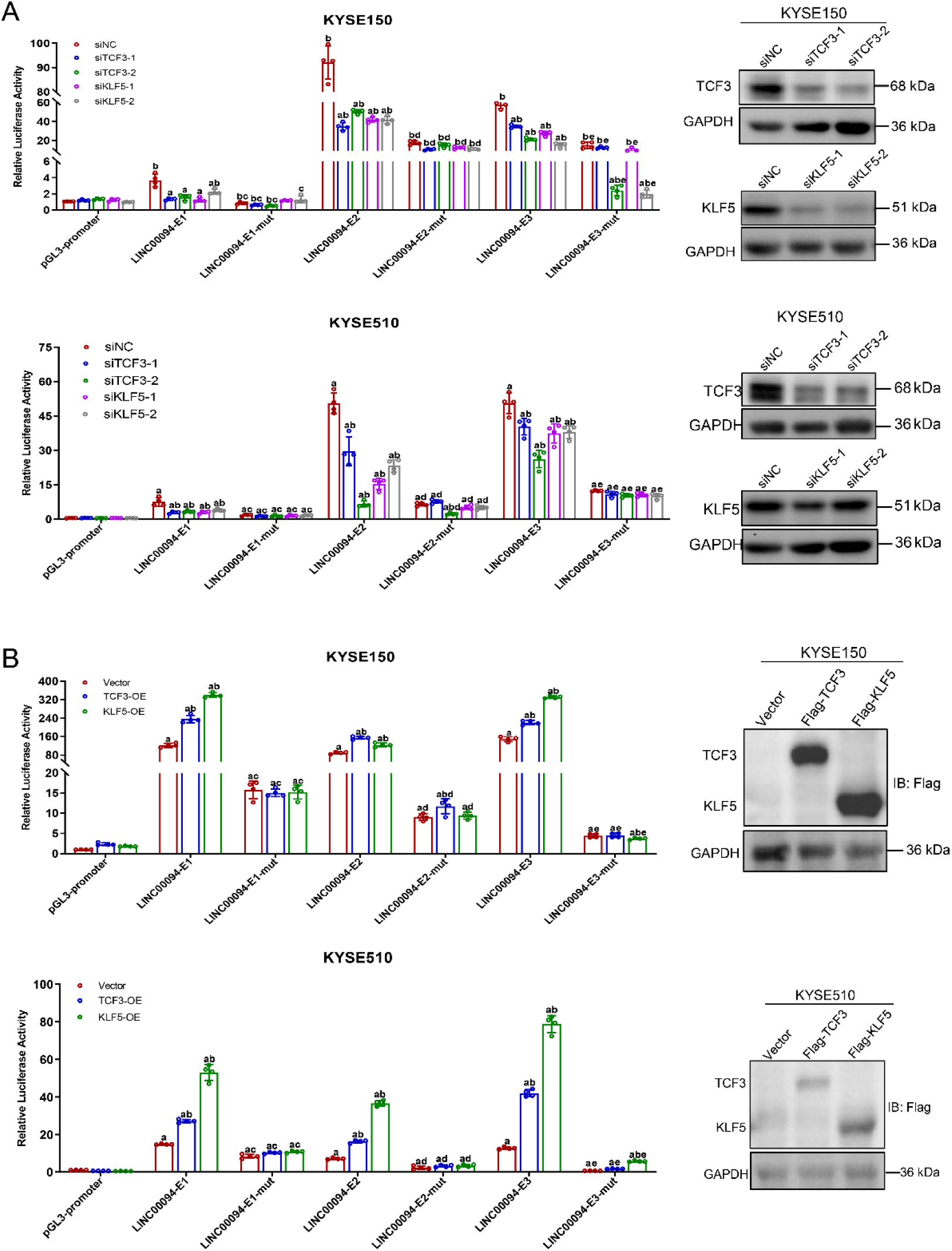
Luciferase reporter assay measuring the activity of LINC00094 SE in KYSE150 and KYSE510 cells. **A.** Left panel, LINC00094 enhancer plasmid, and enhancer mutant plasmid were transfected in KYSE150 and KYSE510 cells after knocking down TCF3 or KLF5. The cells were harvested 48 h later, and the reporter gene activity was measured. The firefly luciferase activity was normalized to Renilla luciferase activity, and the relative value from the cells transfected with the pGL3-promoter was set to 1. Right panel, the expression of TCF3 and KLF5 were evaluated in the KYSE150 cells using western blotting. **B.** As described in **A**, the luciferase reporter gene assay after overexpression of TCF3 and KLF5. Each value represents the mean ± SD, n≥3. a: siRNA or overexpression group compared with siNC or vector group, *P* < 0.05; b: pGL3-promoter-enhancer or pGL3-promoter-enhancer-mutant compared with pGL3-promoter-vector, *P* < 0.05; c: E1-mutant compared with E1, *P* < 0.05; d: E2-mutant compared with E2, *P* < 0.05; e: E3-mutant compared with E3, *P* < 0.05.

**Figure S6.**
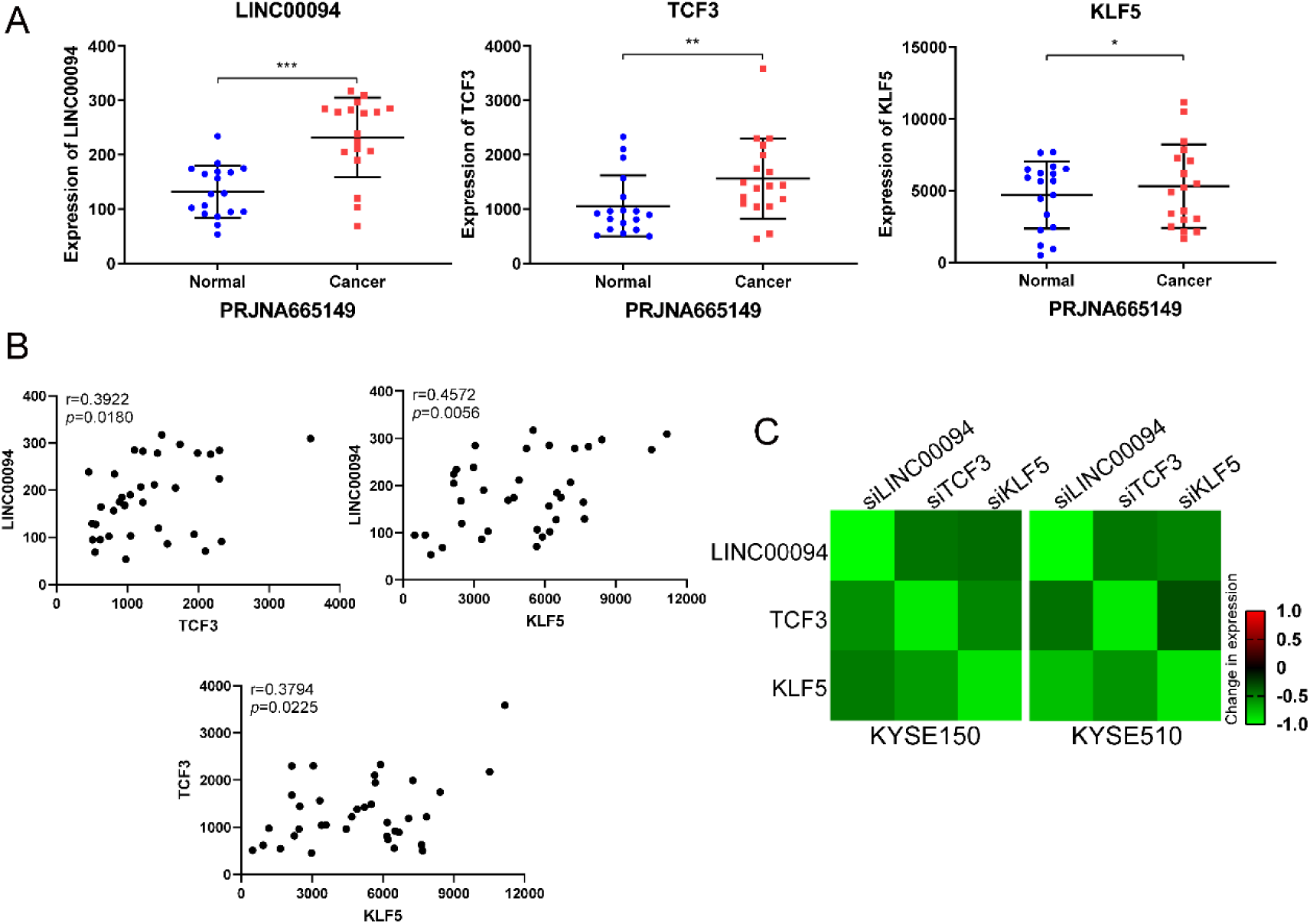
The expression of LINC00094, TCF3 and KLF5 was positively correlated. Related to Figure 4. **A.** Expression of LINC00094, TCF3 and KLF5 in the PRJNA665149 dataset. **B.** Pearson correlation analysis of TCF3, KLF5 and LINC00094 using PRJNA665149 dataset (r>0 positive correlation, r<0 negative correlation). **C.** Heatmap of LINC00094, TCF3, and KLF5 expression fold change as a result of RNA-seq.

**Figure S7.**
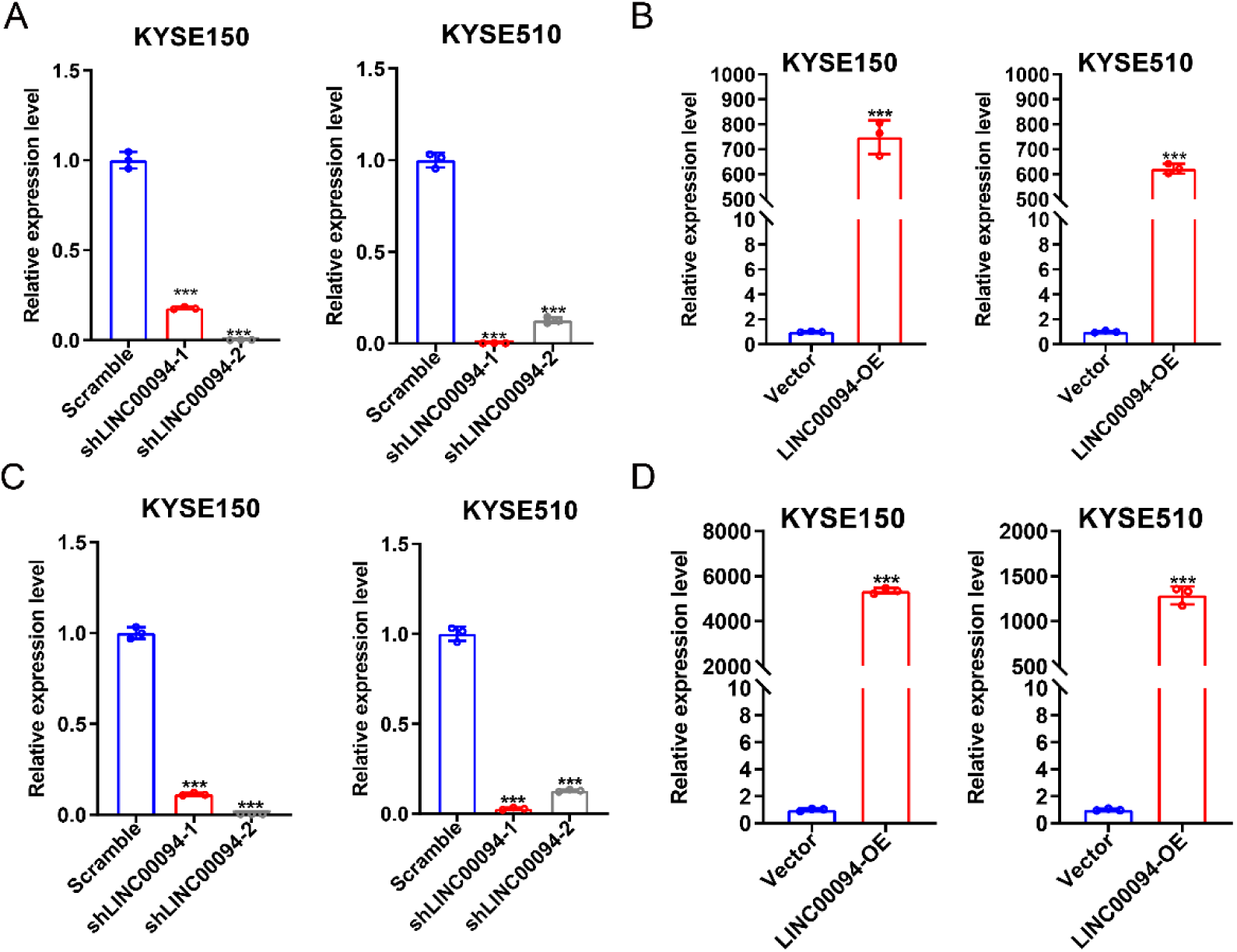
Knockdown and overexpression efficiency of LINC00094. Related to Figure 4. TCF3 and KLF5 expression in KYSE150 and KYSE510 cells detected by qRT-PCR after either LINC00094 knockdown (**A** and **C**) or LINC00094 overexpression (**B** and **D**). **A** corresponds to the result of figure 4E; **B** corresponds to the result of figure 4F; **C** corresponds to the result of figure 4F; **D** corresponds to the result of figure 4G. Each value represents the mean ± SD, n≥3. *** *P* < 0.001. *P*-values were determined using a two-sided t-test.

**Figure S8.**
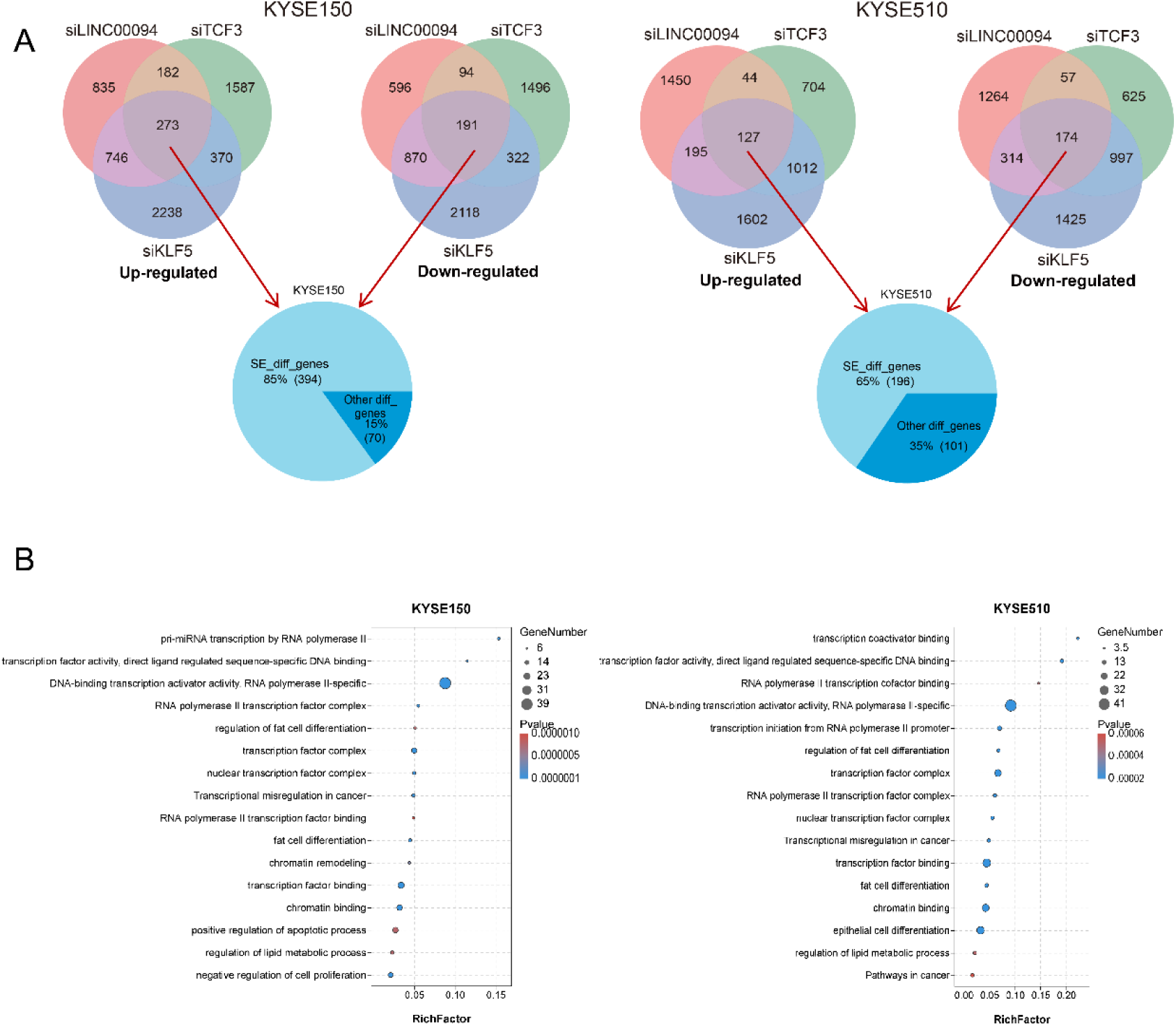
LINC00094, TCF3, and KLF5 co-regulate lipid metabolism pathway in ESCC. **A.** The most common DEGs related to TCF3, KLF5, and LINC00094 knockdown are regulated by SEs. 85% (394/464) the common DEGs knocked down by LINC00094, TCF3 and KLF5 were regulated by SEs in KYSE150 cells, whereas 65% (196/301) were regulated by SEs in KYSE510 cells. **B.** GO enrichment analysis of commonly downregulated genes upon LINC00094, TCF3, and KLF5 knockdown. Color of circles denotes fold changes, and dot size represents the number of genes enriched.

**Figure S9.**
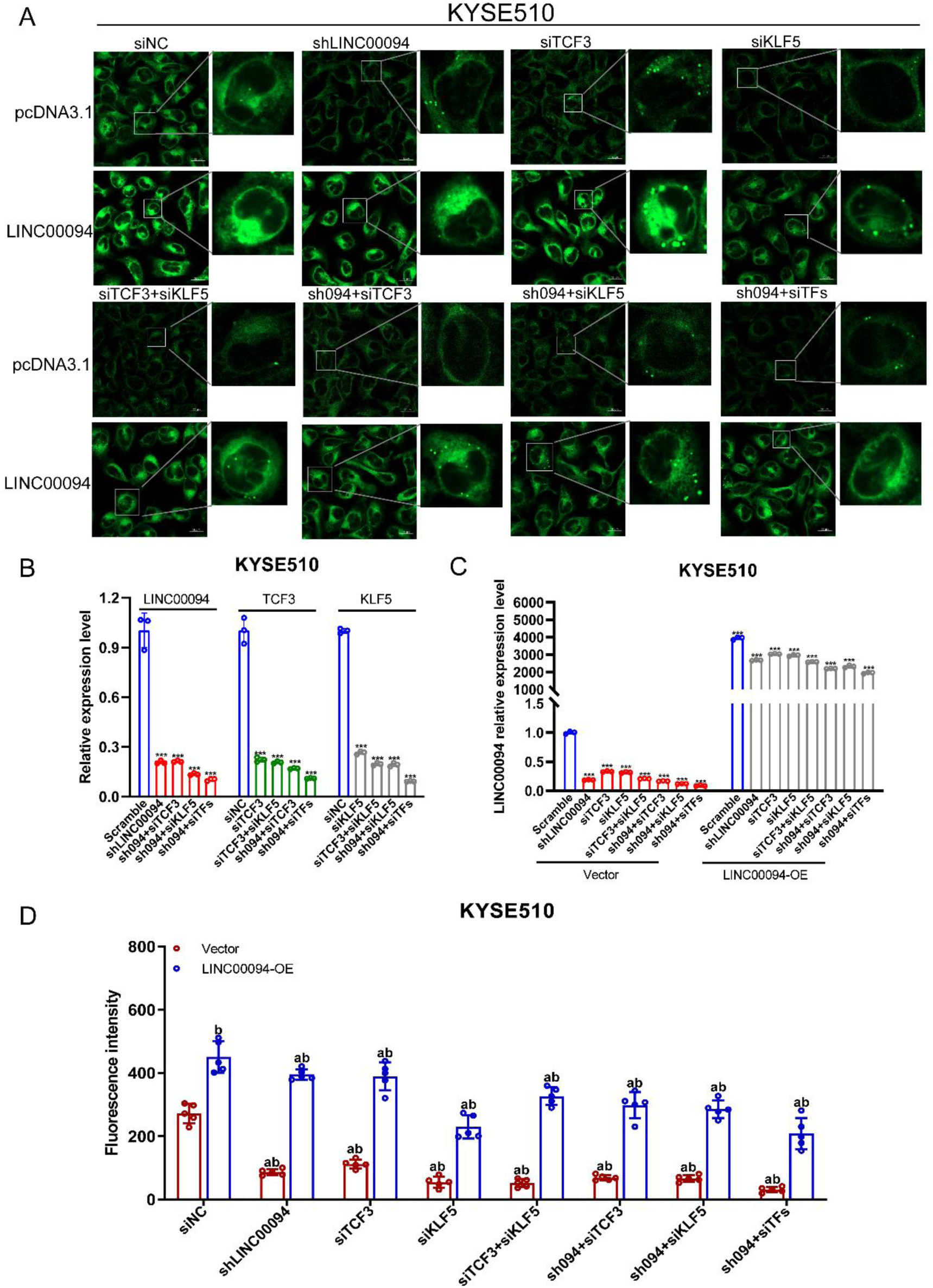
LINC00094 rescues lipid droplet reduction caused by silencing of LINC00094, TCF3, and KLF5 in KYSE510 cells. Related to Figure 6. **A.** Confocal images of staining of lipid droplets after either LINC00094/TCF3/KLF5 knockdown or combined with full-length LINC00094 overexpression in KYSE510 cells (sh094: shLINC00094; siTFs: siTCF3+siKLF5; 094-OE: LINC00094 overexpression). Scale bar, 20 μm. **B-C.** qRT-PCR analysis was performed to detect the mRNA levels of LINC00094, TCF3, and KLF5 after knockdown (**B**) or overexpression (**C**) in different treatment groups of lipid droplet staining assay. Each value represents the mean ± SD, n≥3. * *P* < 0.05, ** *P* < 0.01, *** *P* < 0.001. **D.** Quantitative analysis of lipid droplet staining based on the confocal images; Mean ± SD are shown, n = 5, as the number of microscopic vision. a: siRNA or overexpression group compared with siNC or vector group, *P* < 0.01; b: LINC00094-OE compared with vector, *P* < 0.01.

**Figure S10.**
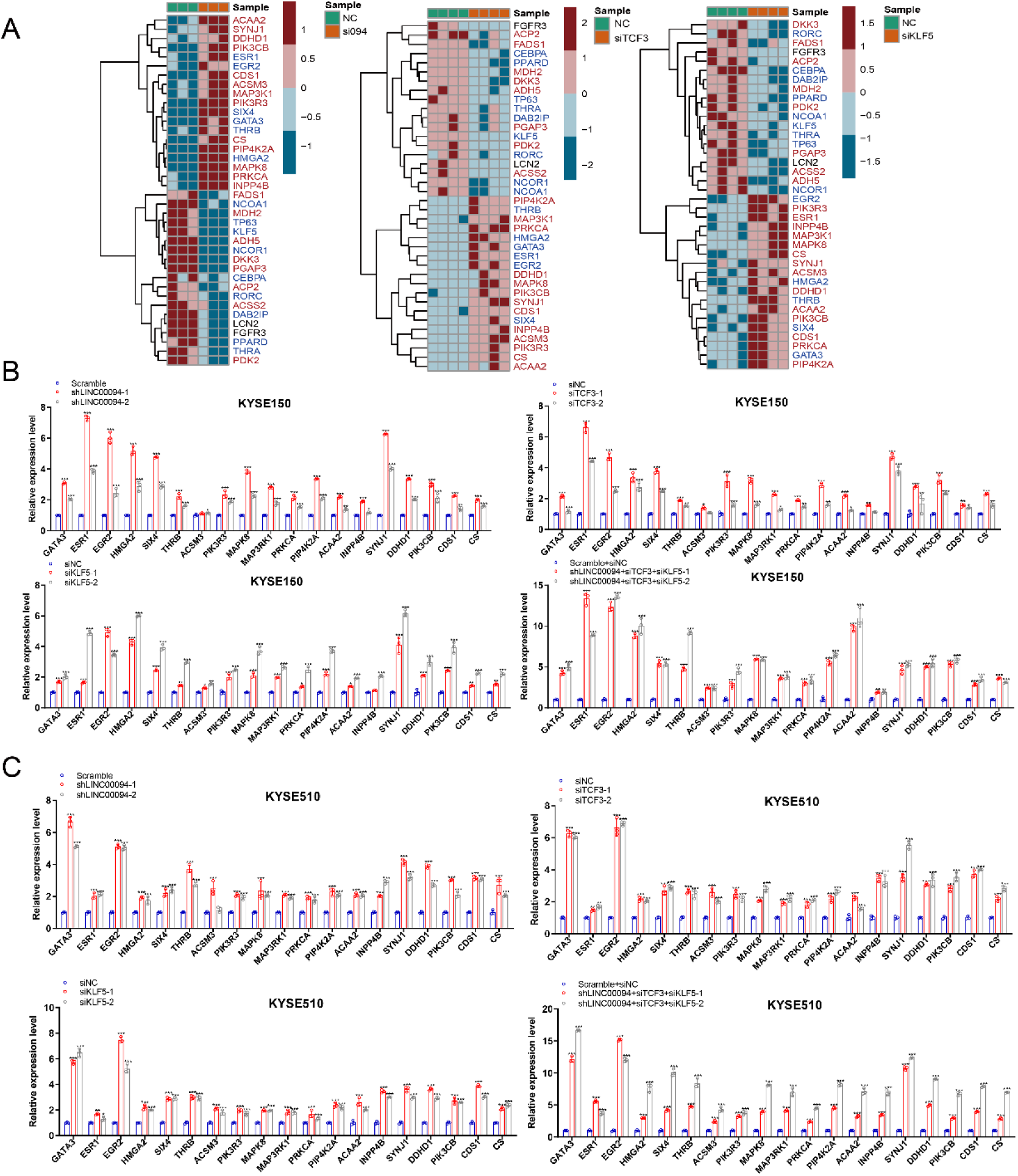
LINC00094, TCF3, and KLF5 regulate lipid metabolism-related genes expression. Related to Figure 6. **A.** Cluster analysis was performed on the common differential genes after knocking down LINC00094, TCF3, and KLF5. Red represents key enzymes in lipid metabolism, and blue represents transcription factors. **B-C.** mRNA level of lipid metabolism-related genes after knocking down LINC00094, TCF3, and KLF5 in KYSE150 (**B**) and KYSE510 (**C**) cells. Each value represents the mean ± SD, n≥3. * *P* < 0.05, ** *P* < 0.01, *** *P* < 0.001. P-values were determined using a two-sided t-test.

**Figure S11.**
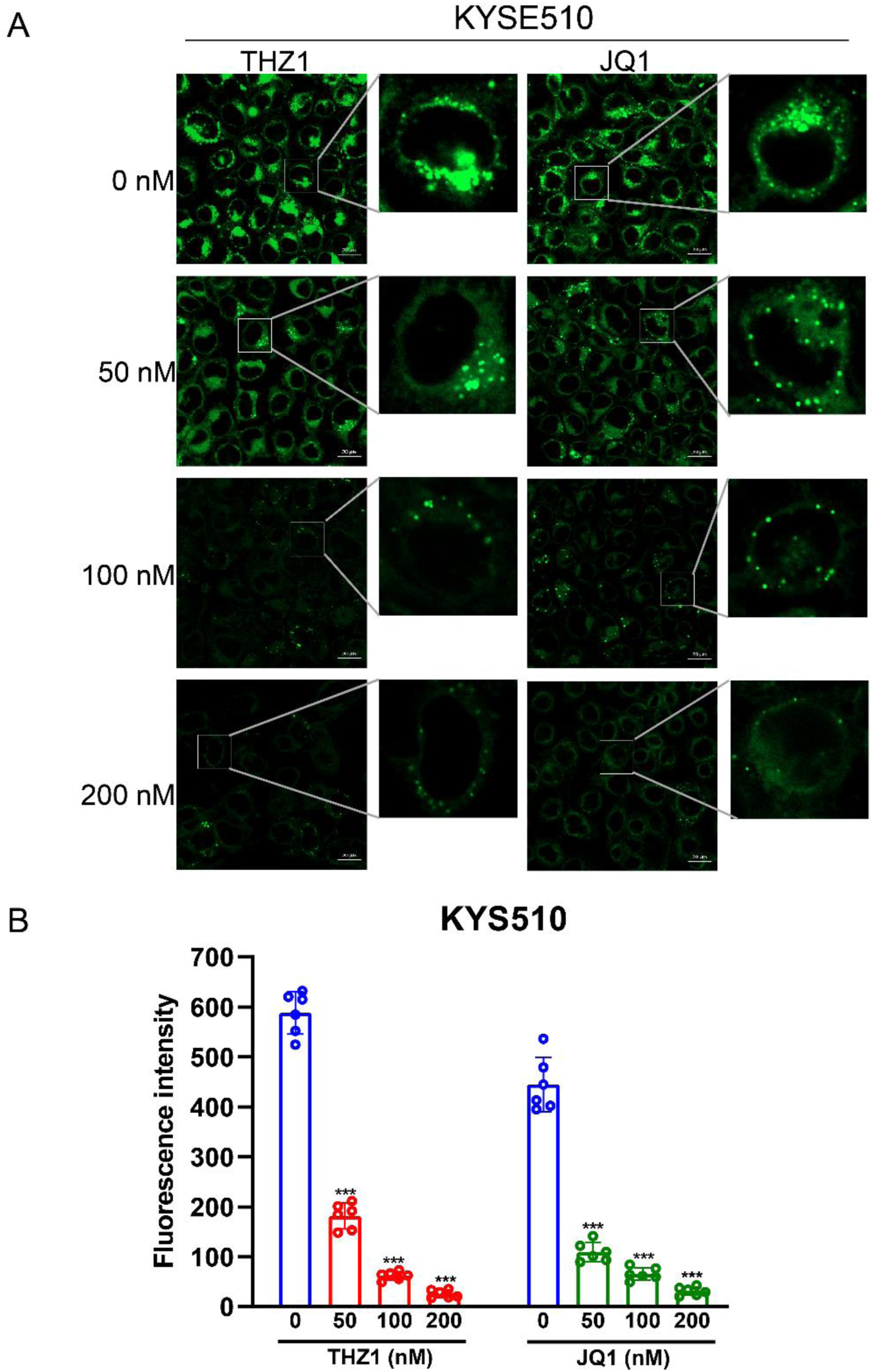
THZ1 and JQ1 treatment effectively inhibited the formation of lipid droplets in KYSE510 cells. Related to Figure 7. **A.** Confocal images of staining of lipid droplets after THZ1 and JQ1 treatment (0, 50, 100, 200 nM, 12 h) in KYSE510 cells. **B.** Quantitative analysis of lipid droplet staining based on the confocal images; Mean ± SD are shown, n = 6, as the number of microscopic vision. *** *P* < 0.001. *P*-values were determined using a two-sided t-test.

**Figure S12.**
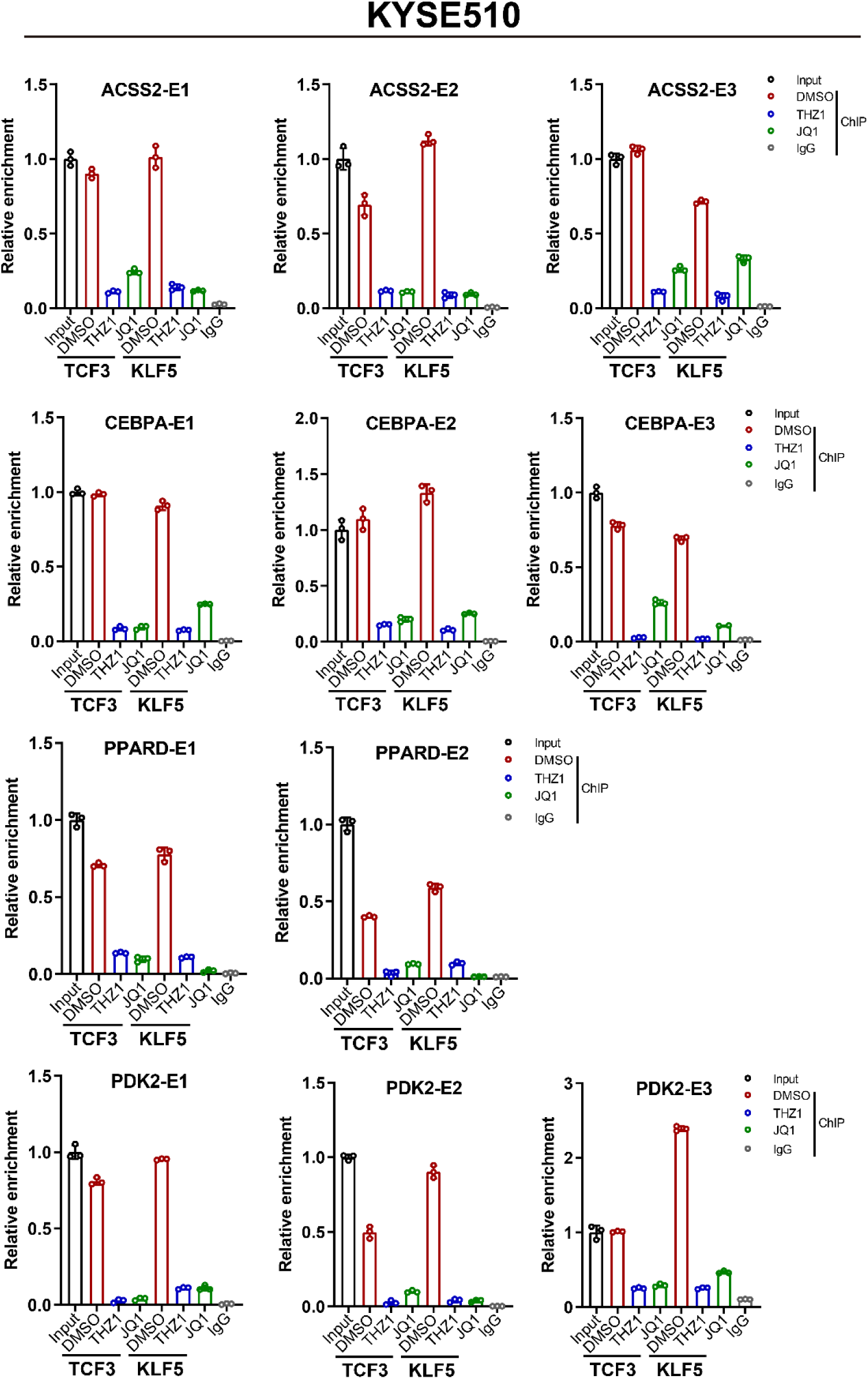
TCF3 and KLF5 bind directly to lipid metabolism-related gene SEs to regulate their expression. Related to Figure 7. ChIP-qPCR experiments measuring TCF3 and KLF5 binding on the ACSS2/PDK2/CEBPA/PPARD SE segments upon treatment with THZ1 (100 nM,12 h) and JQ1 (100 nM, 24 h) in KYSE510 cells. The technical triplicates in a representative experiment are shown and performed twice. Error bars indicate the mean ± SD from three replicates per group. IgG represents the NC antibody.

**Figure S13.**
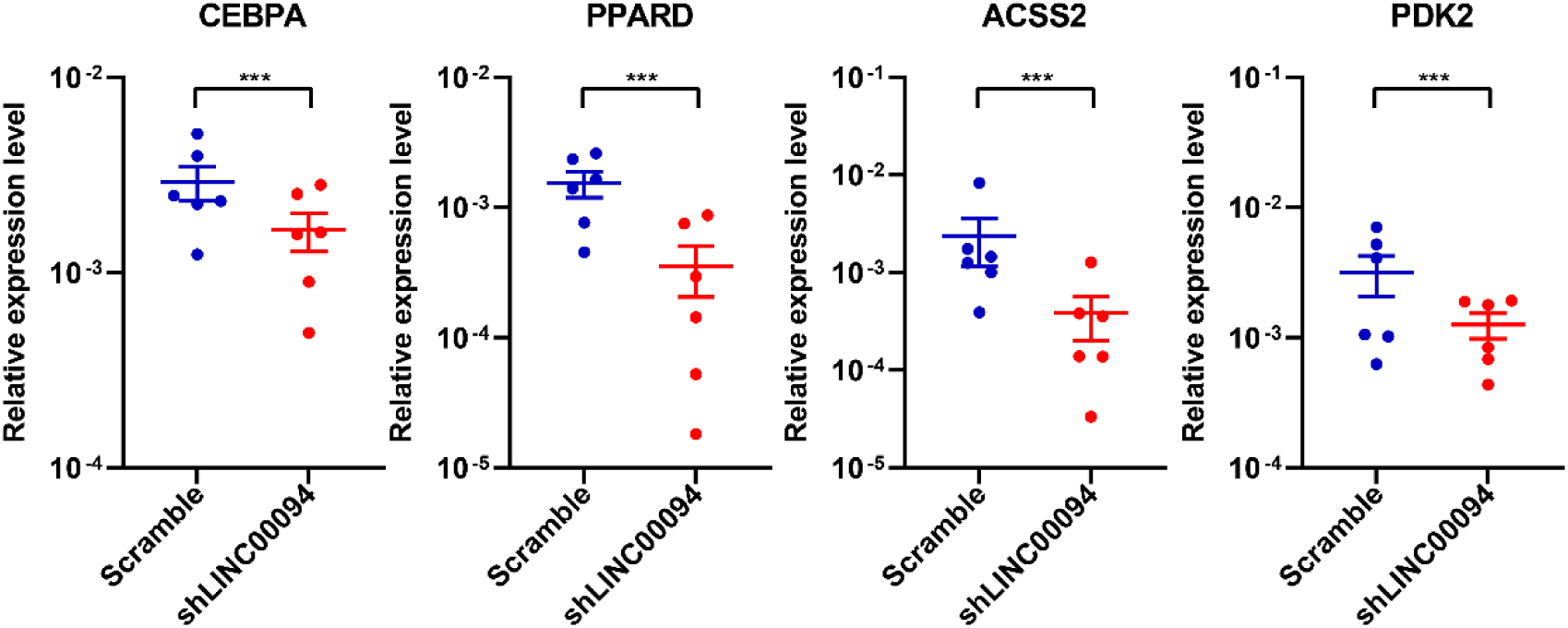
LINC00094, TCF3, and KLF5 promote lipid synthesis. qRT-PCR assay to detect mRNA levels of lipid metabolism-related genes in xenograft mouse tumors. The mean ± SEM are shown, n = 6, *** *P* < 0.001. *P*-values were determined using a two-sided t-test.

**Figure S14.**
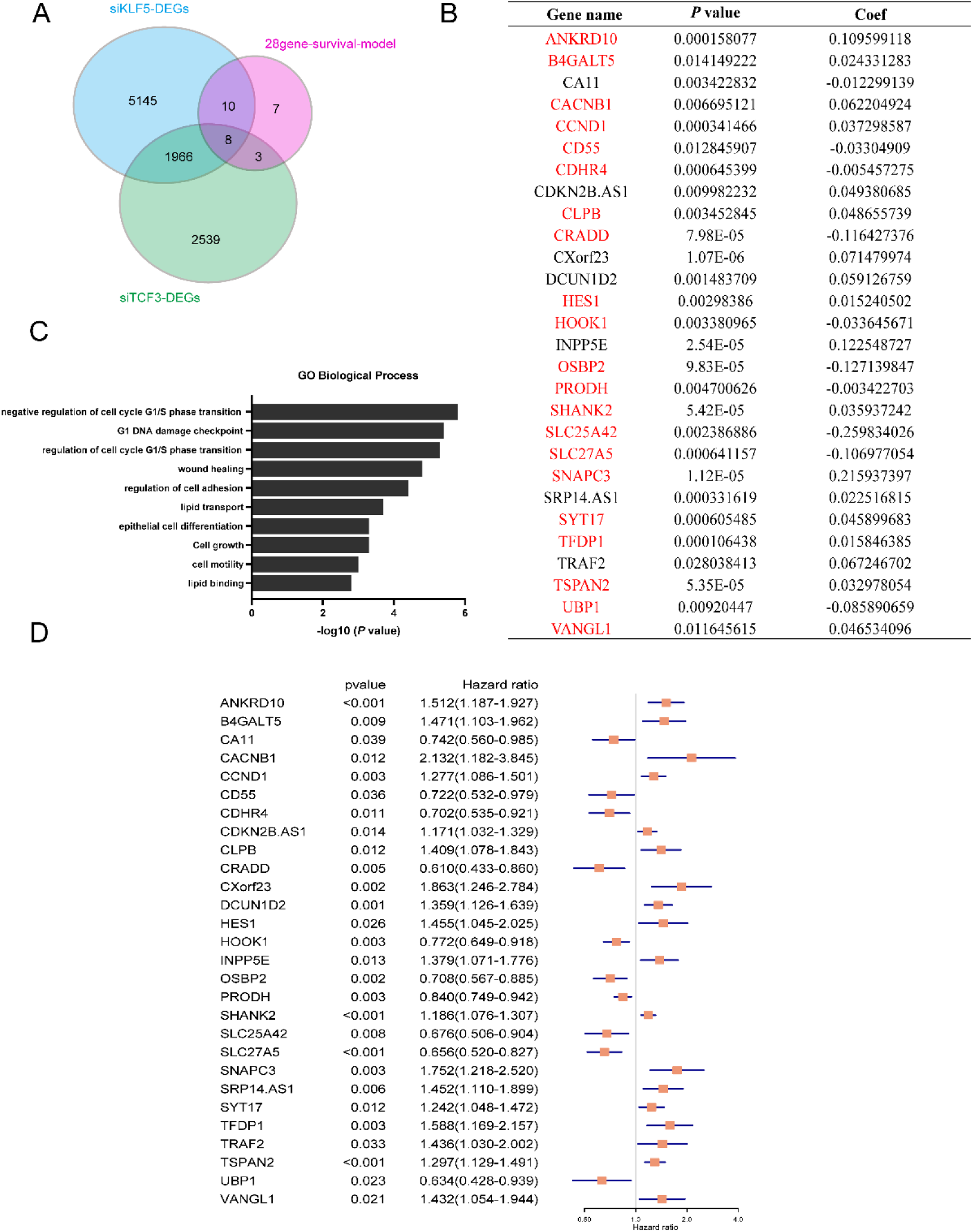
Most genes of the LASSO prognostic signature are co-regulated by LINC00094, TCF3, and KLF5. **A.** Venny map and the prognostic signature of DEGs after knocking down TCF3 and KLF5. **B.** The list constituted 28 genes representing the prognostic signature, *P* < 0.05 represents meaningful survival, and coef represents the LASSO regression coefficient. Red font indicates genes regulated by TCF3, KLF5, and LINC00094. **C.** GO enrichment analysis of commonly regulated genes in the prognostic signature. **D.** Univariate Cox analyses for these 28 genes.

**Figure S15.**
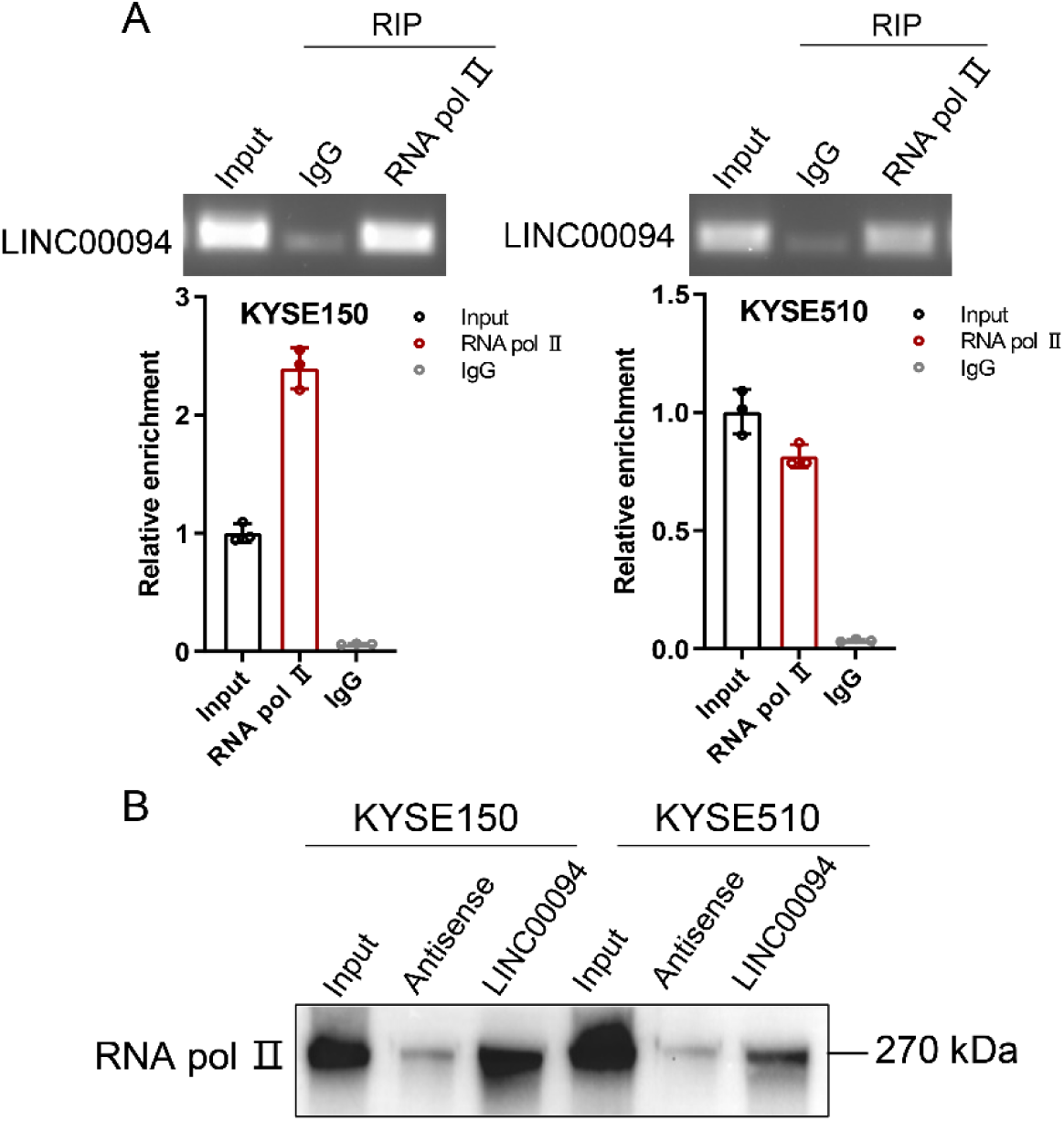
The ternary complex of LINC00094, TCF3, and KLF5 bind to RNA pol Ⅱ. **A.** RIP followed by RT-PCR was performed to detect the interaction between RNA pol Ⅱ and LINC00094 in KYSE150 and KYSE510 cells. Each value represents the mean ± SD, n≥3. **B.** RNA-pulldown assay of biotin-labeled full-length LINC00094 RNA in KYSE150 and KYSE510 cells. Western blotting of RNA pol Ⅱ in the LINC00094 complex.

**Supplementary Table S1.**
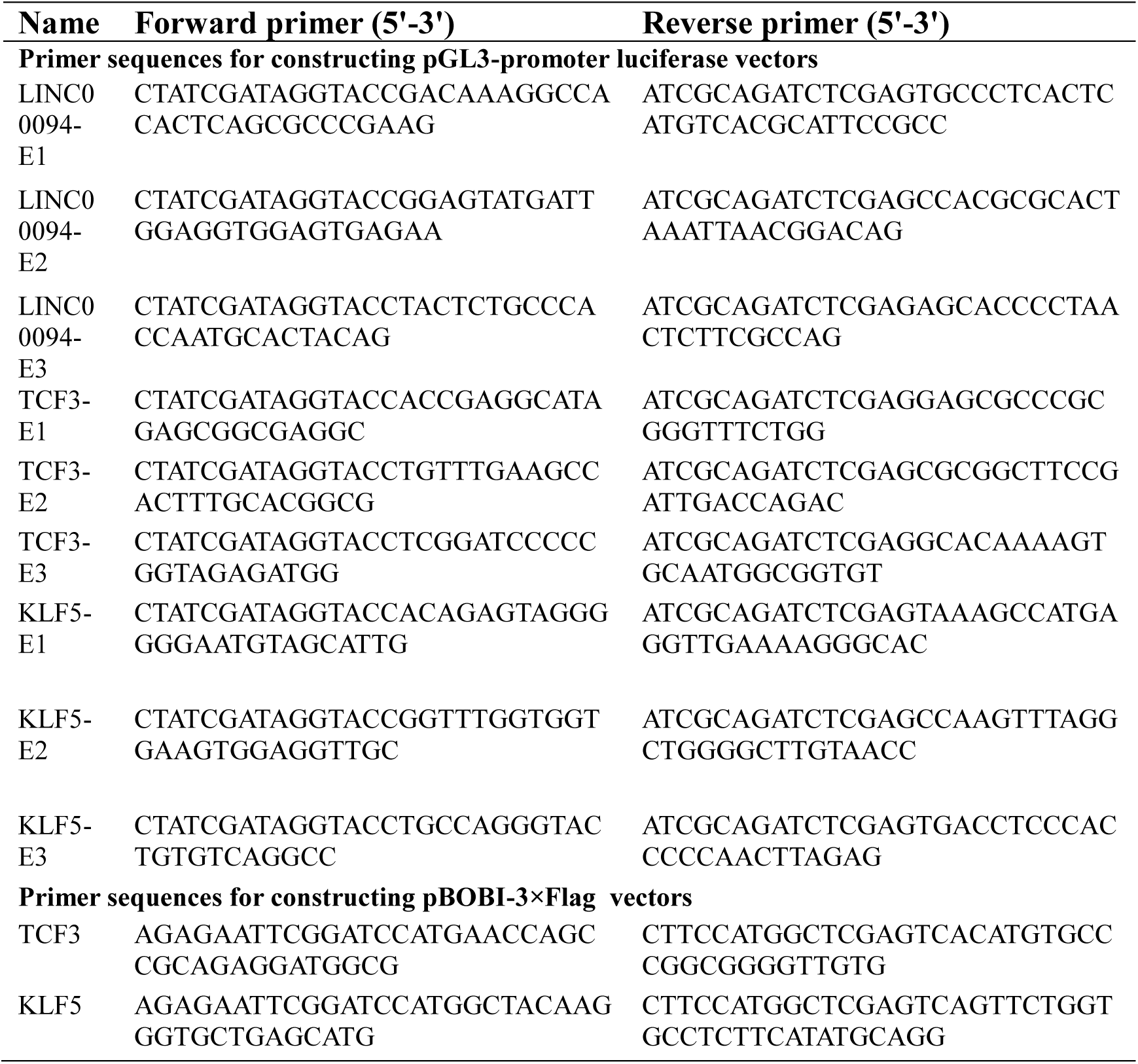
Primer sequences for gene clone.

**Supplementary Table S2.**
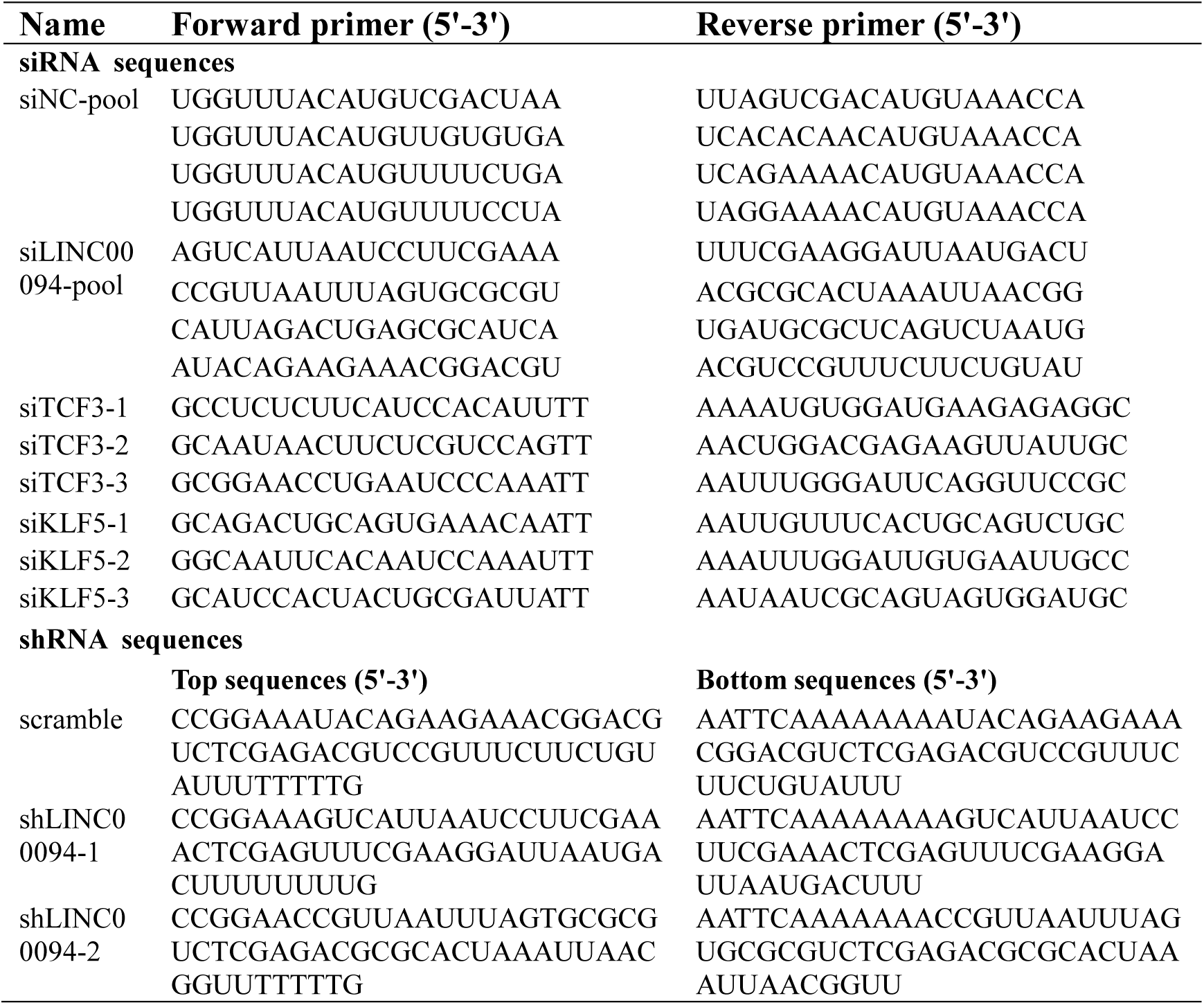
The siRNA and shRNA target sequences.

**Supplementary Table S3.**
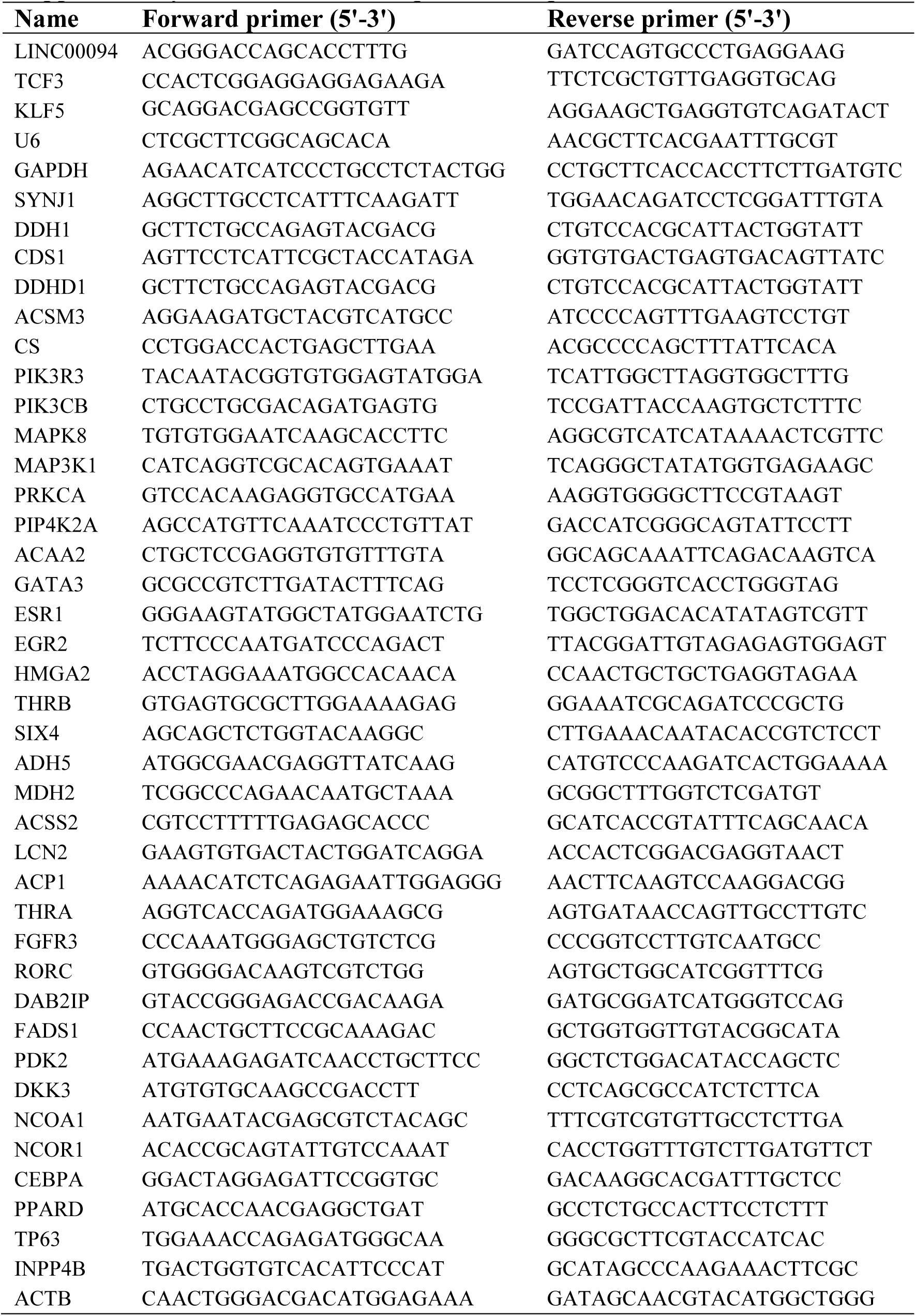
Primer sequences for quantitative RT-PCR.

**Supplementary Table S4.**
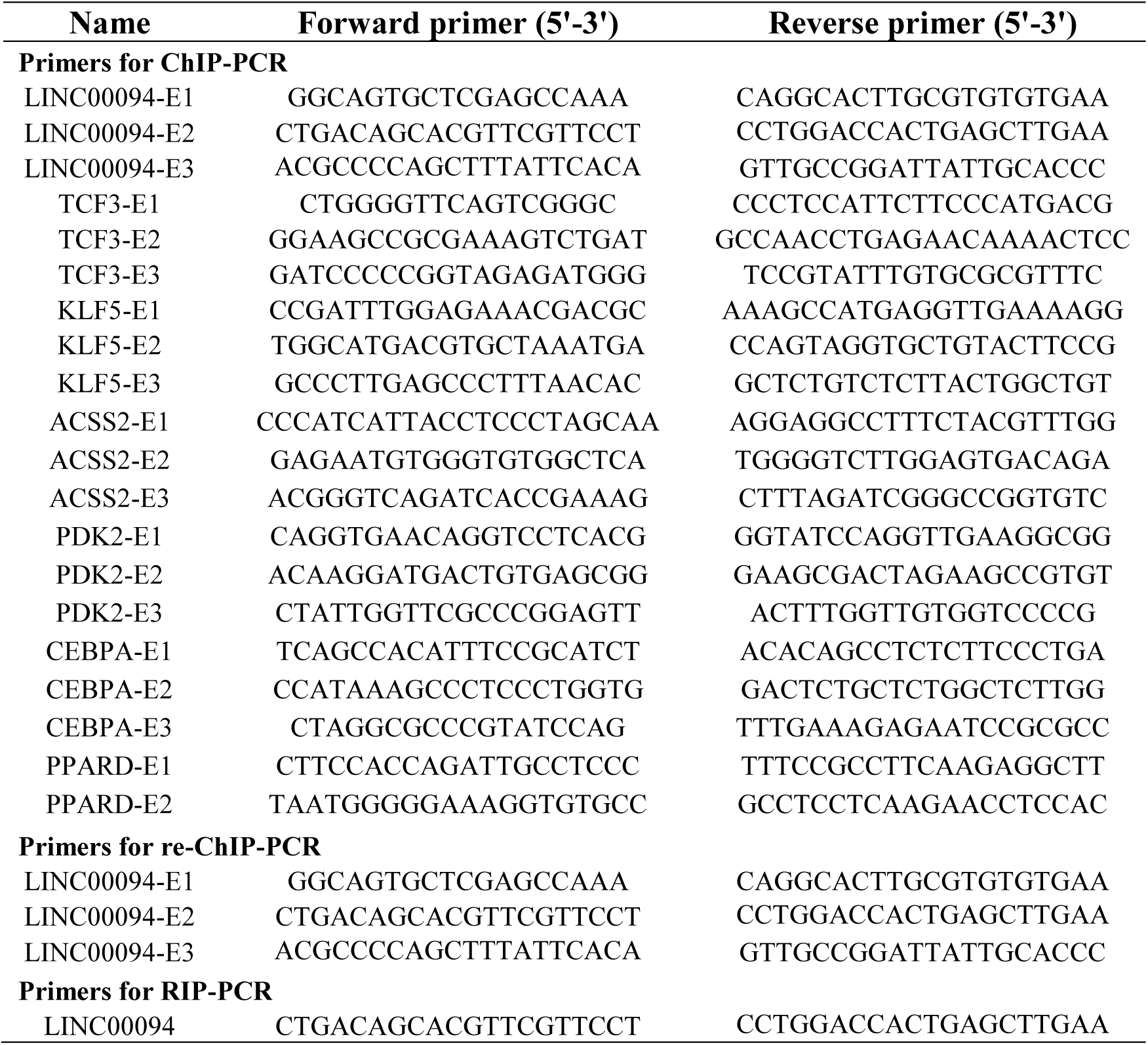
Primer sequences for ChIP-PCR, re-ChIP and RIP.

